# Satellite glial translatome reprograms to favor mitochondrial and vesicular signaling under pelvic pain conditions

**DOI:** 10.64898/2026.11.11.766000

**Authors:** Sathish Kumar Yesupatham, Alyssa Granley, Alison Xiaoqiao Xie

## Abstract

Sensory satellite glial cells actively contribute to inflammatory and neuropathic pain, but their roles in pelvic pain remain unclear. This study examined how the active translating mRNAs in sensory SGCs change during pelvic pain symptoms using translating ribosome affinity purification followed by RNA sequencing (TRAP-seq) in a chronic pelvic pain (CPP) mouse model. Our data showed that SGCs exhibit sex-specific translatome patterns at baseline, which were reprogrammed in CPP conditions under the control of eukaryotic translation initiation factor 4E (*Eif4e*). In CPP conditions, sensory SGCs exhibit increased translation of mitochondrial and vesicular genes and suppression of immune genes in both sexes; an anti-inflammatory cytokine profile was detected in female pelvic sensory ganglia. We are the first to characterize SGC translatome in pelvic pain conditions and to report sex differences in SGC translatome. Our data suggested that mitochondrial and vesicular gene translation in sensory SGCs may be important to afferent hypersensitivity.

**Highlights:**

- TRAP-seq resolves the satellite glial cell translatome in mouse lumbosacral DRG.
- Sensory SGC translatome exhibits sex-specific signatures at baseline.
- Immune-related gene translation in SGCs is suppressed during pelvic pain.
- SGCs increase mitochondrial and vesicular gene translation in CPP condition.

## Introduction

Chronic pelvic pain (CPP) is highly prevalent worldwide and imposes substantial personal and economic burden^1,2^. Many CPP patients experience urinary frequency, urgency, and voiding difficulties resembling overactive bladder (OAB). Vascular endothelial growth factor (VEGF) has been identified as a urine biomarker for CPP ^3,4^; intravesical VEGF instillations in mice induce afferent sprouting, OAB symptoms, and pelvic mechanical hypersensitivity similar to those observed in CPP patients^5^, providing a mouse model of CPP for mechanistic study.

Recently, we performed RNA-seq profiling in lumbosacral dorsal root ganglia (LS-DRG), where the pelvic sensory neuronal somata reside. Our data identified sex-specific transcriptional changes in mouse LS-DRG following VEGF instillations, suggesting that different immune and glial cell types were involved in developing pelvic hypersensitivity in each sex^6^. In addition, distinct contributions of immune cells and spinal microglia to pain have been reported in males and females^7,8^. In this study, we decided to focus on satellite glial cells (SGCs) in LS-DRG and asked how SGC translatome changes during CPP.

SGCs envelope individual sensory neuronal soma, regulate the perineuronal environment^9,10^, and undergo changes that can modulate neuronal excitability and immune signaling following nerve injury^11,12^. However, the molecular programs of SGCs in pain states remain poorly defined. Their flattened morphology and tight neuronal association complicate isolation^13^, and culture rapidly alters their phenotype and transcriptome^14^. Although single-cell (sc) and single-nucleus (sn) atlases have resolved SGC identity and heterogeneity^15–17^, their ability to detect low-abundance transcripts is limited by assay sensitivity and sequencing depth^18,19^. In this study, we performed translating ribosome affinity purification followed by RNA sequencing (TRAP-seq) in sensory SGCs surrounding pelvic sensory neurons for glial-specific, sex-stratified integration of CPP mechanisms. TRAP-seq captures ribosome-associated mRNA from genetically defined cells directly from homogenized tissue, without cell dissociation^20–22^. It has been applied to CNS cell types and DRG nociceptors^23–27^.

Our data revealed sex-specific translatome landscape in sensory SGCs at the baseline, while CPP symptoms were associated with changes in selected trafficking and mitochondrial transcripts. Immune-related transcripts were significantly changed without evidence of broad cytokine activation in the LS-DRG. These data provide a resource and testable hypotheses for SGC translational regulation in visceral pain.

## Results

### Sox10-guided ribosomal GFP expression in LS-DRG was restricted to SGCs and limited numbers of Schwann cells

SGCs and Schwann cells (myelinating and non-myelinating) represent the majority of the DRG cells expressing *Sox10* gene^28^. Low levels of *Sox10* expression was also detected in small populations of DRG neurons and endothelial cells^28^. To verify which cell population expresses ribosomal GFP tag in our study, TRAP mice (*Sox10-CreER^T2+/−^::L10a-GFP^+/f^*) underwent tamoxifen induction and their LS-DRG were isolated for confocal imaging of GFP signals (Fig. 1A-B). The TRAP mice are expected to have OFP expression in Sox10+ cells without tamoxifen; tamoxifen induction is expected to cease the OFP expression while inducing L10a-GFP expression. Therefore, the ratio of GFP+/OFP+ cells were used to estimate recombination efficiency. In TRAP mice injected with tamoxifen, we observed both GFP+ and OFP+ cells in the ratio of 3:2 (Fig. 1B), suggesting 60% transduction rate following the current tamoxifen protocol (Fig. 1C). Among all the GFP+ cells, 77.25 ± 3.48% were positive for SGC marker BLBP and 3.52 ± 2.03% were positive for Schwann cell marker MPZ, suggesting that majority of the L10a-GFP+ cells were SGCs (Fig. 1D). In TRAP mice injected with corn oil, only 3 GFP+ cells were detected (Fig. 1B). No OFP+ or GFP+ cells were observed in *Cre-*animals with either tamoxifen (6 DRG/2 animals) or corn oil injections (7 DRG/2 animals) (data not shown). These data demonstrate the specificity and efficiency of the TRAP model for profiling the SGC translatome. All experimental animals in the study were tamoxifen-induced TRAP mice that underwent intravesical instillations of VEGF_165_ or saline.

**Figure 1.**
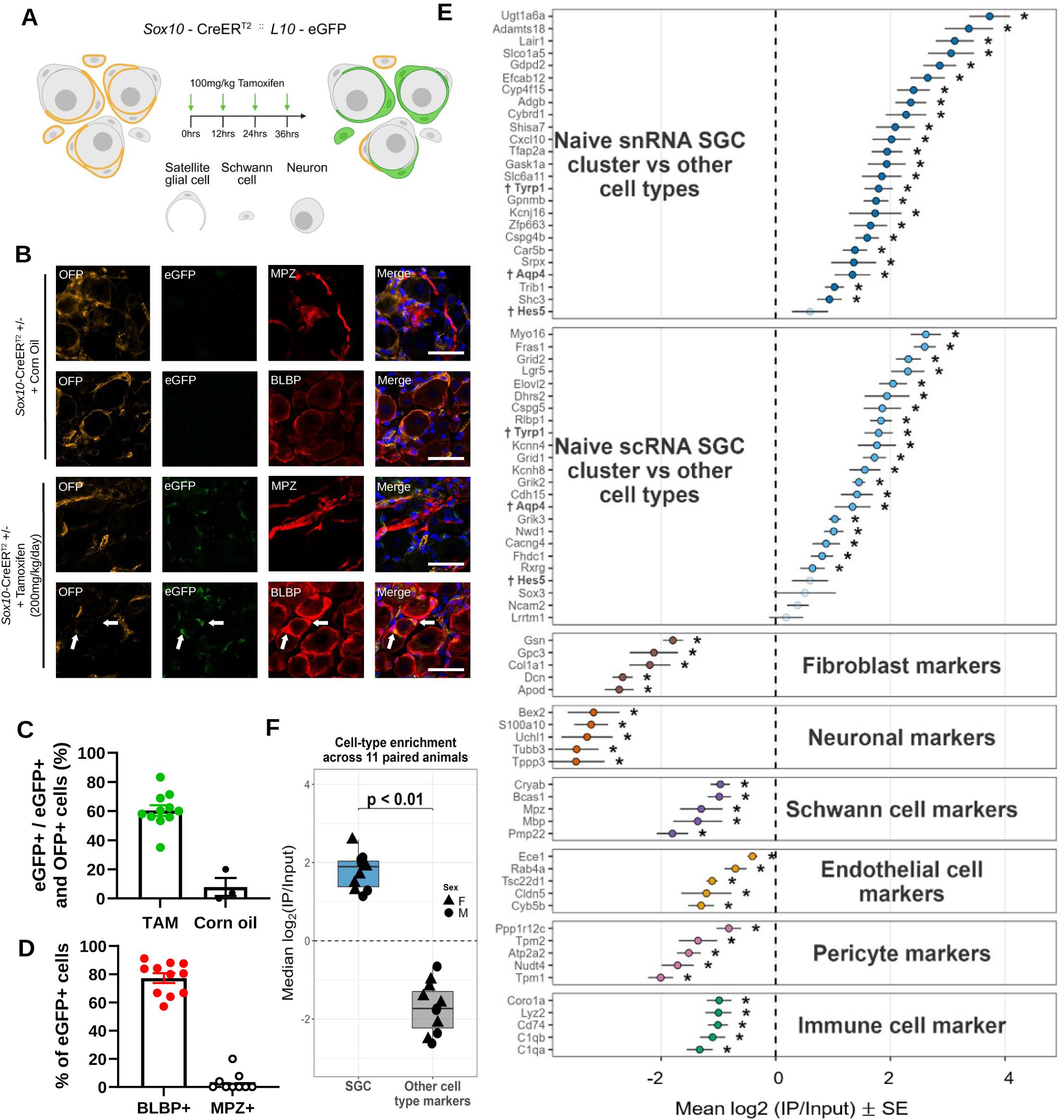
Sox10-TRAP enriches the SGC translatome and depletes other DRG cell-type transcripts. (A) Tamoxifen injections induce L10a-GFP expression in SGCs. (B) OFP and GFP expression in DRG from TRAP mice. White arrows point to BLBP+ SGCs. Scale bar=40µm. (C) Percentage of eGFP+ cells out of total eGFP+ and OFP+ cells suggests ∼60% transduction rate in the DRG following the current Tamoxifen induction protocol. Each dot represents one confocal image. (D) GFP+ cells in DRG are largely BLBP+. Each dot represents one image; the bars are SE. (E) Per-gene enrichment of cell-type marker panels in TRAP-IP versus subject-matched input evaluates SGC translatome enrichment. Each point represents the mean log₂(IP/input) ± SE. Per-gene significance was tested using two-sided paired Wilcoxon signed-rank tests against zero, followed by Benjamini–Hochberg FDR correction within each marker panel. Bright points and asterisks indicate FDR<0.05; and faded points indicate FDR ≥ 0.05. The vertical dashed line marks log₂FC = 0. The top two panels compare the TRAP-IP data with SGC-enriched genes identified in the snRNA and scRNA atlases. Overlapping genes from both datasets are in bold and dagger-marked (†). The lower panels compare TRAP-IP data with non-SGC enriched genes in fibroblast, neuron, Schwann cell, endothelial cell, pericyte, and immune cells. (F) Cell-specific enrichment summary across 11 paired animals. 47 SGC enriched genes and 30 non-SGC genes were included, and the enrichment score was calculated as the median log₂(IP/Input) across the snRNA and scRNA SGC marker genes. A non-SGC score was calculated as the mean of class medians across the six non-SGC marker panels, giving each cell class equal weight. SGC and Non-SGC scores were compared using a two-sided paired Wilcoxon signed-rank test. Boxes show median and IQR; points represent individual animals. Circles indicate males and triangles indicate females. The significance bracket shows the paired Wilcoxon result.

### Intravesical instillations VEGF induced pelvic hypersensitivity in both male and female mice

Repeated intravesical instillations of a mouse VEGF recombinant protein, VEGF_165_ results in low bladder capacity, urinary frequency, and pelvic mechanical hypersensitivity^29^. Intravesical instillations of VEGF_165_ in *Sox10-CreER^T2^* mice induced significant increases in withdrawal frequencies to Von Frey filaments (Supplemental Fig. 1A, N=3 in each sex), which lasted until week 8 (Supplemental Fig. 1B), suggesting that this model is suitable for studying CPP mechanisms. To study SGC translatome, both sexes of *Sox10-CreER^T2^::L10a-GFP* mice were subjected to intravesical instillations of VEGF_165_ (pelvic pain group, males, N=3; females, N=3) or saline (control group, males, N=3; females, N=3) (Supplemental Fig. 1C). On Day 16, the LS-DRG (L1-L2, L6-S2) were collected, and the translatome patterns were compared between sex and between the CPP and control groups.

### The enrichment of SGC translatome was validated using cell-specific gene panels derived from independent DRG sc and sn RNA-seq atlases

To verify whether the immunoprecipitated ribosome-bound RNA (TRAP-IP) enriched SGC translatome, the TRAP-IP data from individual animals were compared with their own Input (total RNA from the same DRG), creating subject-matched IP-Input pairs (N=11, 5 males and 6 females). The top 25 genes expressed in sensory SGCs were identified from two independent, naive DRG SGC clusters^14,30^, providing 50 top-expressed SGC genes (STAR methods) used in Figure 1E. In the TRAP-IP data, these 50 genes were largely enriched when compared to the Input, as evidenced by the positive mean log₂(IP/Input) values and Benjamini–Hochberg (BH)-adjusted FDR<0.05 (Fig. 1E). SGC cluster annotation was concordant between the two atlases, 629 were shared between the scRNA (963 genes) and snRNA (1,074 genes). Importantly, cell type-specific genes for fibroblasts, neurons, Schwann cells, endothelial cells, pericytes, and immune cells (five per cell type; Supplementary Fig. S2) based on the atlas^28^ were consistently depleted in the TRAP-IP fraction, suggesting that TRAP-IP enriched translatome in largely SGCs. We also performed cell-type enrichment analysis using 47 SGC-enriched genes and 30 non-SGC genes (Fig. 1F). On average, the SGC marker panel was significantly enriched in the TRAP-IP fraction (∼2 median log2 IP/Input) compared to the non-SGC marker panel (−1.5 median log2 IP/Input) across all animals (two-sided paired Wilcoxon, p<0.01, N=11). The observed SGC gene enrichment is consistent between sexes and instillation groups. Together, these findings demonstrate that the TRAP-seq data selectively captures SGC mRNAs in LS-DRG.

### TRAP-IP revealed three main categories of genes under active translation in sensory SGCs

We next asked which genes are under active translation in sensory SGCs. Sensory SGCs showed robust translation of gap junction and GPCR signaling related genes, represented by TRAP-IP counts per million (CPM) in Figure 2 (left column). The Input CPM is shown in the middle column; high CPM in TRAP-IP and low Input CPM suggest enrichment in SGCs. SGC pseudobulk CPM calculated from a naïve DRG snRNA-seq atlas^30^ was used to compare available transcripts with translating mRNA levels (Fig. 2, right column). Differential expression of a particular gene between TRAP-IP and SGC pseudobulk suggests translational regulation. Our analysis revealed active translation in sensory SGCs of genes associated with gap junctions and GPCR signaling (Fig. 2A and D), while housekeeping genes were underrepresented in TRAP-IP relative to the Input (Fig. 2G). Specifically, *Gja1* (Cx43) showed the highest expression (47.8 CPM) with concordant pseudobulk expression (95.4 CPM), suggesting high transcription and translation levels of *Cx43* in sensory SGCs (Fig. 2A-C). *Gjc1* (Cx45), *Panx1* (Panx1), and *Panx2* (Panx2) showed high CPM in TRAP-IP but not in SGC pseudobulk RNA data, suggesting that these gap junction genes might be actively translated from relatively low numbers of transcripts in sensory SGCs.

**Figure 2.**
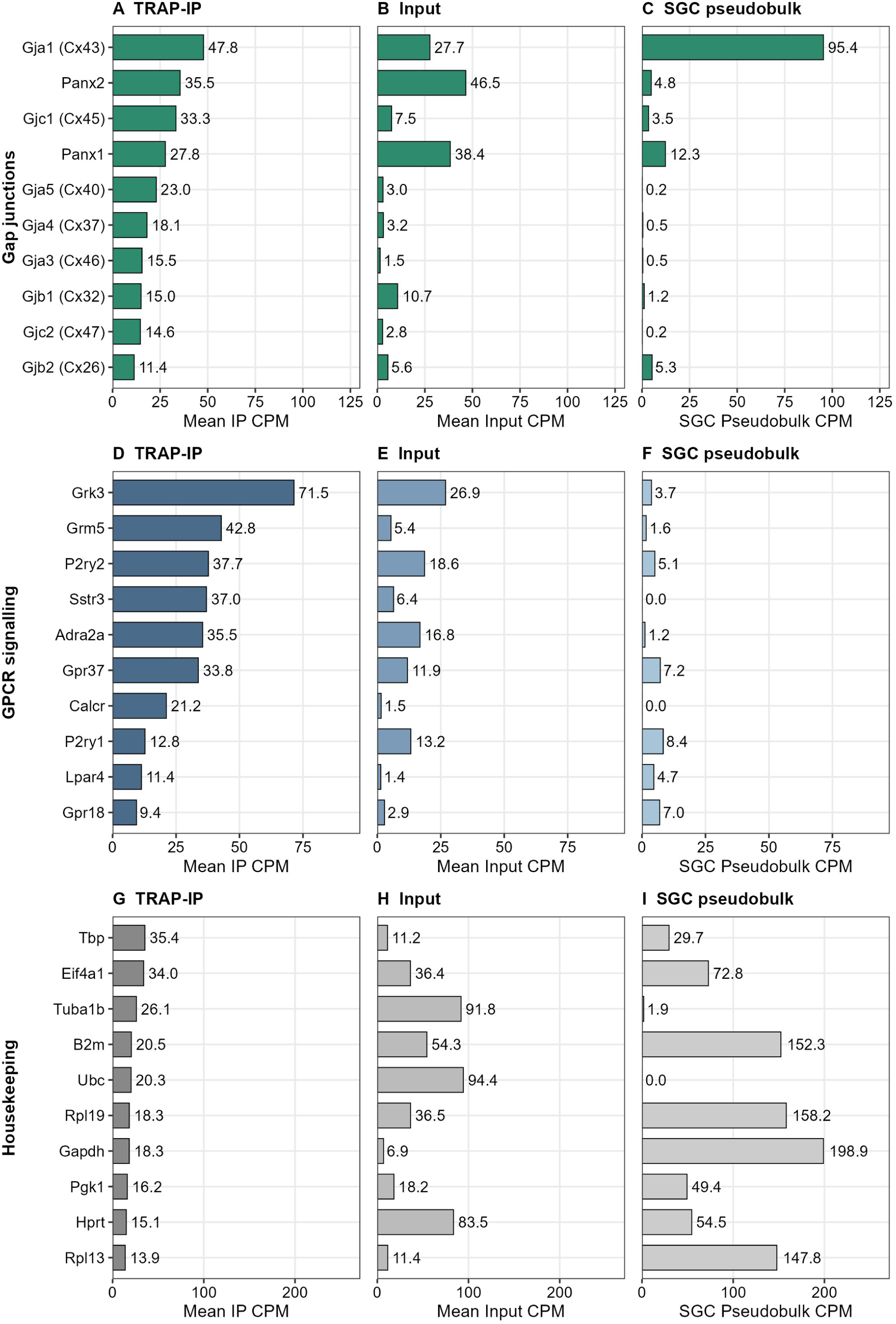
Sox10-TRAP captures SGC signaling transcripts across gap junction and GPCR pathways but not housekeeping genes. (A–C) Expression of selected connexin and pannexin family transcripts in TRAP-IP RNA (A), bulk DRG input RNA (B), and SGC pseudobulkRNA from existing snRNA-seq data (C). TRAP-IP robustly captures SGC-associated gap junction genes including *Gja1 (Cx43)*, *Panx2*, *Gjc1 (Cx45)*, *Panx1*, and *Gjb1 (Cx32)*. In contrast, several connexin/pannexin transcripts show weak representation in SGC pseudobulk despite clear enrichment in TRAP-IP samples. (D–F) Expression of SGC-associated GPCR signaling transcripts in TRAP-IP RNA (D), bulk DRG input RNA (E), and SGC pseudobulk RNA (F). TRAP-IP samples show strong representation of signaling-related genes including *Grk3*, *Grm5*, *P2ry2*, *Sstr3*, *Adra2a*, and *Gpr37*, whereas several of these transcripts are detected at lower levels in the input and substantially low SGC pseudobulk data. (G–I) Housekeeping gene controls in TRAP-IP RNA (G), bulk DRG input RNA (H), and SGC pseudobulk snRNA-seq data (I). Housekeeping values reflect uniquely mapped reads only; genes with many processed pseudogenes (e.g., Gapdh) are underestimated under unique-read counting, and housekeeping genes are shown for reference rather than as normalisation controls. Bars represent mean counts per million (CPM) across 11 matched TRAP-IP and bulk DRG input samples. SGC pseudobulk CPM values represent the mean of male and female naïve DRG SGC pseudobulk snRNA-seq datasets.

Among the GPCR signaling genes, *Grk3* (GPCR kinase 3), *Grm5* metabotropic glutamate receptor 5), *P2ry2* (P2Y2 purinergic receptor), *Sstr3* (somatostatin receptor type 3), *Adra2a* (alpha-2A adrenergic receptor), *Gpr37* (G protein-coupled receptor 37), and *Calcr* (calcitonin receptor) are the most abundant in SGC translatome (Fig. 2D), suggesting their importance in DRG neuron-SGC communication. However, SGC pseudobulk CPM of these genes were much lower and even when compared to the Input CPM, suggesting strong positive translational regulation (Fig. 2D-F).

Housekeeping genes, including *Gapdh* (glyceraldehyde 3-phosphate dehydrogenase), *Rpl19* and *Rpl13* (60S ribosomal protein subunits), and *B2m* (beta-2-microglobulin), were among the most abundant transcripts in SGC pseudobulk data (Fig. 2I) but were underrepresented in the TRAP-IP fraction (Fig. 2G) in relative to Input (Fig. 2H). Because TRAP counts were restricted to uniquely mapped reads, genes with many processed pseudogenes, such as *Gapdh*, are underestimated. These differences therefore should not be interpreted as differences between transcription and translation.

### Satellite glia express a GPCR repertoire dominated by contact and positional receptors

We next examined GPCR genes in the SGC translatome using the IUPHAR/BPS Guide to Pharmacology target list and a three-step filter (STAR Methods; Fig. 3A). Of 343 GPCRs evaluated, 119 exceeded the empirical detection floor (24.4 CPM), 40 of these were confirmed in bulk FACS-sorted naïve SGCs^13^, and 32 remained after excluding receptors dominated by non-SGC expression in the two DRG atlases^14,30^. The retained receptors are predominantly adhesion GPCRs (n = 9), Frizzled/Smoothened GPCRs (n = 6), together with lysophospholipid receptors, *Celsr1* and *Celsr2*, *Ednra* and *Ednrb*, *Gprc5b* and *Gprc5c*, *Lgr4* and *Lgr5*, *Adcyap1r1*, *Lhcgr*, *Gabbr1*, *Adora1*, and *Gpr37* and *Gpr37l1* (Fig. 3B). *Gabbr1* was the most abundant receptor in the immunoprecipitated (highest CPM), followed by *Celsr2*, *Gprc5b*, *Celsr1*, *Fzd3* and *Lpar1*. Aminergic and chemokine/complement receptors that cleared the detection floor were excluded at the atlas step because they were dominated by non-SGC expression. Next, TRAP-IP abundance of the 32 GPCRs was compared with paired whole-DRG Input (Fig. 3C). The relative expression level spans approximately −1 to +3.4 log₂ units. Position on either axis reflects how much of a transcript is present in the translatome rather than whether a receptor is expressed: *Gpr37l1* sits near the detection floor in Figure 3B and near zero in Figure 3C, despite broad expression across SGCs in both atlases^14,30^.

**Figure 3.**
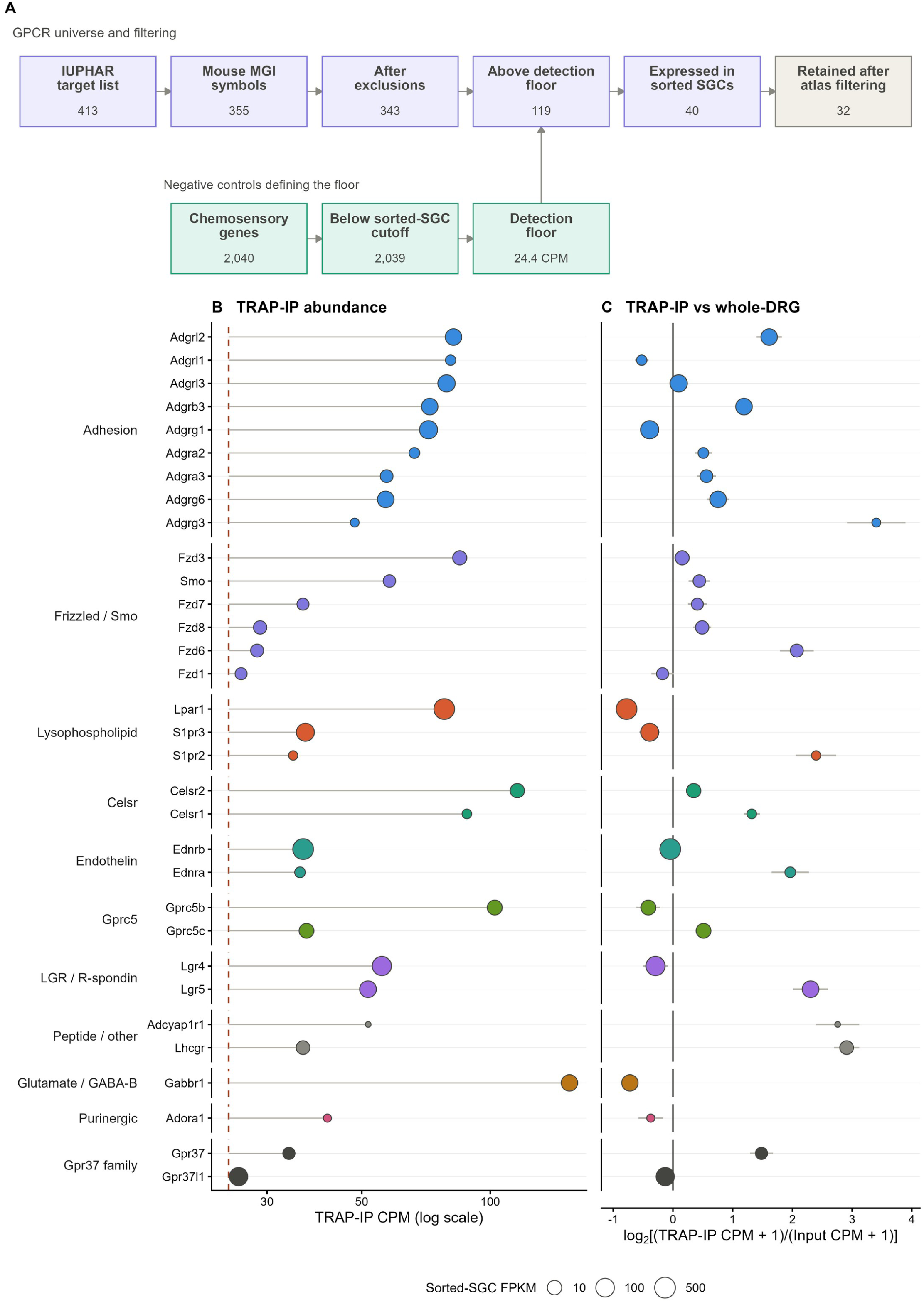
Sox10-TRAP reveals a 32-receptor candidate SGC repertoire. (A) GPCR filtering workflow. The IUPHAR/BPS list contained 413 GPCR-type entries representing 355 distinct mouse genes. The list was reduced to 343 receptors after excluding IUPHAR-designated non-GPCR 7TM proteins, *Adgrd2-ps*, *Gpr31c*, chemosensory receptors, and genes absent from the translatome count matrix. Presumed chemosensory negative controls that did not meet the sorted-SGC expression criterion (2,039 genes) were used to define the detection floor as their 95th percentile mean TRAP-IP abundance (24.4 CPM). Of the 343 receptors, 119 exceeded this floor and 40 met the sorted-SGC criterion of FPKM >1 in at least three of four naïve replicates. Eight were excluded because cell-weighted non-SGC expression was at least 2.5-fold higher than SGC expression in each of the two DRG atlases, leaving 32 candidates. Box values show gene or receptor counts. (B) Mean abundance of the 32 receptors across eleven TRAP-IP libraries (five saline and six VEGF) grouped by receptor family. The x-axis represents TRAP-IP CPM on a logarithmic scale, and the horizontal line running from the dashed floor shows abundance above the detection floor. Point size represents mean expression in sorted naïve SGCs. (C) Mean TRAP-IP-to-input ratio calculated within eleven matched animal pairs (five saline and six VEGF). Positive values indicate greater representation in TRAP-IP, whereas negative values indicate greater representation in whole-DRG input. Error bars show the standard error across pairs. These values describe relative RNA representation, not receptor activation.

In a sensitivity analysis restricted to the five saline pairs (STAR Methods), the detection floor was 25.5 CPM, and the selection yielded 33 receptors: 31 shared with the pooled repertoire, with *Fzd1* present only in the pooled set and *Gpr146* and *Gpr173* only in the saline-only set (Supplementary Fig. S3). All three lie within 4 CPM of the floor, which is estimated less precisely from five libraries than from eleven. The pooled repertoire was retained as the better-powered estimate.

### Baseline SGC translatome suggested sex-specific SGC signaling

Bulk-RNAseq profiling showed sex-specific gene and pathway changes in LS-DRG associated with VEGF-induced visceral hypersensitivity^6^. Here, we asked whether SGC translatome also exhibits sex differences at the baseline or in CPP conditions. First, the TRAP-IP from saline-instilled mice (N=3 each sex) were compared between sexes (Fig. 4, left column). In saline-instilled animals, 2,229 genes were sex-differentially translated in sensory SGC (p<0.05 and >1.5-fold change) (Supplemental Table S1). Among these 2229 genes, 2,186 (98.1%) were detected exclusively in the TRAP-IP, but not in the Input (Fig. 4A), suggesting that these genes are sex-differentially translated in sensory SGCs but likely not in other DRG cell types. Among the 2229 sex-differentially translated genes in sensory SGCs, 768 are autosomal protein-coding genes, 282 are sex-chromosomal (X/Y linked) genes, and 1,179 are non-coding genes (Fig. 4B, all biotypes). The top translationally upregulated genes in male sensory SGCs fall into two distinct programs: chromatin remodeling, represented by the Y-linked histone demethylase *Kdm5d* and the LIM-homeodomain transcription factor *Lhx3*^31,32^, and cytoskeletal organization, represented by the mitotic kinesin motors *Kif18a* and *Kif20a*^33,34^, suggesting coordinated epigenetic and structural regulation in male SGCs. Similarly, the top translationally upregulated genes in female sensory SGCs reflect two functional programs: neuron-glial structural connectivity, represented by the neuronal migration factor *Dcx* and the cell-adhesion molecule *Pcdh19*^35,36^, and transcriptional regulation, represented by the Mediator complex subunit *Med12*^37^, suggesting that female SGCs are translationally primed for neuron-glial communication. This data suggests that male and female sensory SGCs prioritize gene translation for different cell functions.

**Figure 4.**
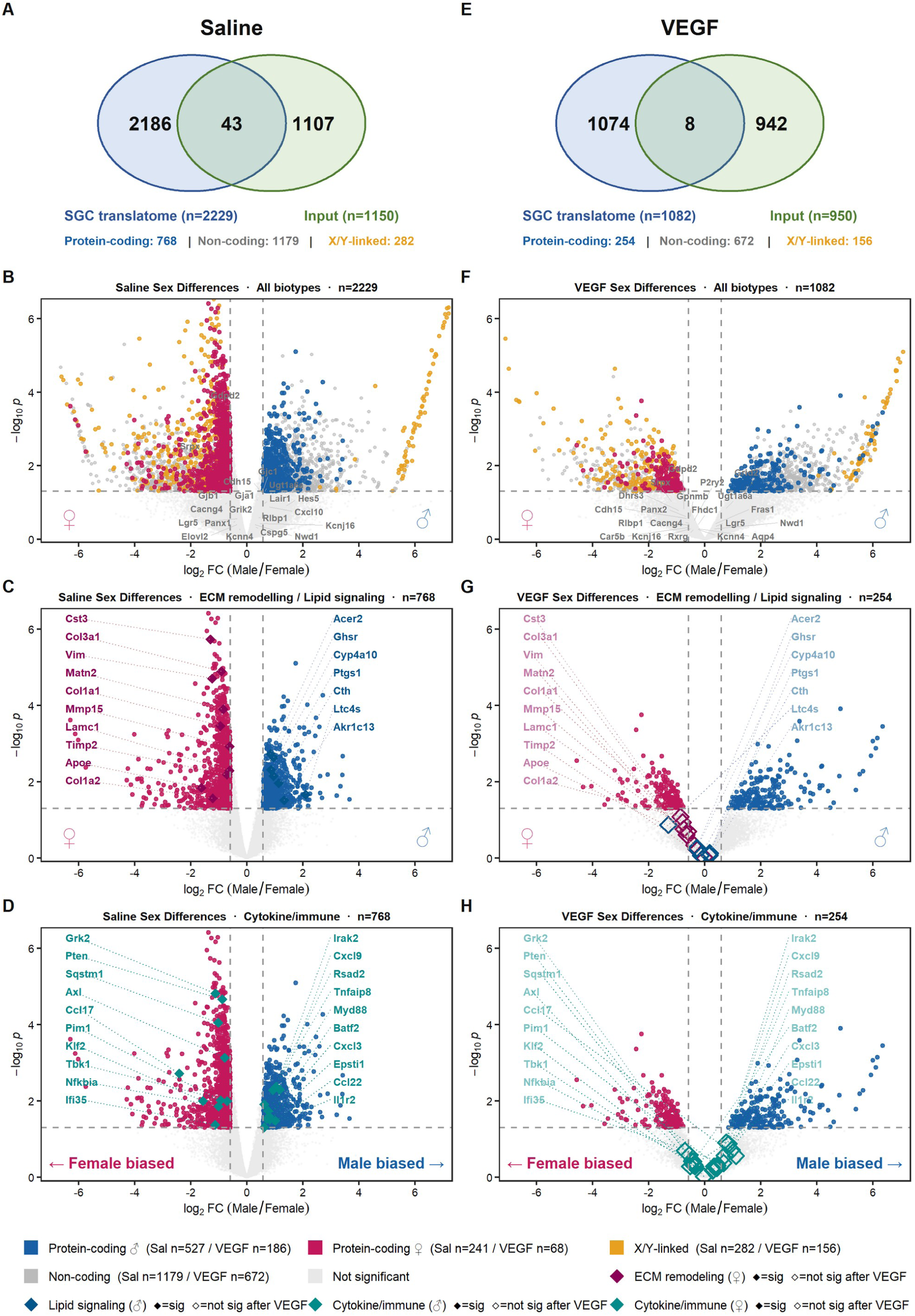
Sex differences in the satellite glial cell (SGC) translatome under saline and VEGF conditions. Male-versus-female differential expression in LS-DRG, shown for saline (A–D) and VEGF (E–H). Differentially expressed genes (DEGs) were defined as nominal p < 0.05 and |log₂FC| > log₂(1.5) (DESeq2 Wald test; model ∼ batch + sex). P-values were not adjusted for multiple testing, so DEG counts are exploratory. **(A, E)** Venn diagrams comparing sex DEGs in the SGC translatome (Sox10-TRAP) and matched whole-DRG Input fractions. Numbers indicate genes that were DEGs in the translatome only, in both fractions, or in the Input only. **(B, F)** Volcano plots of all translatome DEGs across gene biotypes. The biotype breakdown beneath each panel gives the number of DEGs that are protein-coding, non-coding or sex-linked (X/Y). **(C, G)** Volcano plots restricted to autosomal protein-coding genes, highlighting extracellular-matrix (ECM) remodeling genes (female-biased) and lipid/eicosanoid signaling genes (male-biased). **(D, H)** The same autosomal protein-coding genes, highlighting cytokine/immune genes. In all volcano plots, the x-axis shows the log₂ fold-change of male relative to female (positive values indicate higher expression in males), and the y-axis shows −log₁₀(nominal p). Points are colored by category: protein-coding male-biased, blue; protein-coding female-biased, crimson; sex-linked, gold; non-coding, grey; not significant, light grey. Representative SGC marker genes are labeled in grey (B, F). In C, D, G and H, highlighted genes are shown as diamonds connected to gene labels by leader lines and colored by category as indicated in the key. Filled diamonds mark genes meeting the DEG threshold in that condition; open diamonds mark genes significant under saline but not under VEGF. The *n* in each panel title is the total number of DEGs displayed in that panel; category-specific counts are given in the key. Group sizes: translatome, n = 6 per condition (3 male, 3 female); Input, n = 5 for saline (2 male, 3 female) and n = 6 for VEGF (3 male, 3 female).

Analysis restricted to autosomal protein-coding genes (527 male-biased and 241 female-biased genes, Supplemental Table S2) identified distinct sex-biased functional programs (Fig. 4C). Female-biased genes (maroon) included extracellular matrix (ECM) components and remodeling genes (*Col1a1, Col1a2, Col3a1, Lamc1, Vim, Apoe, Cst3, Mmp15, Timp2, and Matn2*), whereas lipid and eicosanoid signaling genes (blue) such as *Ptgs1, Ltc4s, Cyp4a10, Acer2, Akr1c13, Cth, and Ghsr* are male-biased. *Lamc1* (laminin-γ1), an essential basement membrane component required for peripheral nerve glial differentiation and axon myelination^38^; *Vim* (vimentin), a class III intermediate filament and established marker of SGCs^39^. Male-biased genes (blue) included *Ltc4s*, which is involved in leukotriene C4 synthesis and enhances P2X₃-mediated pain responses ^40^. Cyp4a10 encodes a cytochrome P450 hydroxylase involved in lipid metabolism^41^. These data suggest that under basal conditions (saline-instilled and pain-free), male and female SGCs may exhibit different functions. In addition, we identified differential expression of immune-related gene translation in male and female sensory SGCs at the baseline (Fig. 4D).

### Sex-specific SGC translatome patterns are absent under pelvic pain condition

We next asked if there is sexual dimorphism in SGC translatome under CPP conditions. For this analysis, first, the TRAP-IP from VEGF-instilled mice (N=3 each sex) were compared between sexes (Fig. 4, right column). VEGF treatment markedly reduced sex-differential translation, with only 1,082 genes remaining significant (Fig. 4E). Of these, 1,074 (99.3%) were exclusive to the TRAP-IP fraction, which is similar to our finding in saline-instilled animals. Volcano plots of all-biotype sex-differential genes confirmed the marked reduction in significant genes following VEGF treatment (Fig. 4F). Of the 43 SGC marker genes (identified in Fig. 1), 40 (93%) remained within the non-significant region across both saline and VEGF conditions, indicating that core SGC identity genes were largely not sexually differentially translated. Within the autosomal protein-coding subset of genes, male-biased genes decreased from 527 in saline-instilled animals to 186, whereas female-biased genes decreased from 241 to 68. Analysis on the autosomal protein-coding genes showed that the ECM/lipid sex-biased functional programs were largely absent in CPP conditions (Fig. 4G). Overall, the genes that exhibited significant sex-dependent differences in saline-instilled animals fell below the significance threshold in VEGF-instilled animals. Similarly, the sex differences in the immune-related gene subset were largely absent in VEGF-instilled animals (Fig. 4H), including genes involved in PI3K/AKT/mTOR, IFN-γ, inflammatory response, IL-6/JAK/STAT3, and TNF-α/NF-κB signaling. In summary, at the time point examined in our study (day 16 after the first VEGF instillation, when animals exhibited CPP symptoms), SGC gene translation had shifted away from sex-specific programs.

### CPP-associated SGC translatome pattern shifts: from immune transcripts to mitochondrial and vesicular transcripts

We next asked which genes are actively translated in sensory SGCs while animals experience CPP-like pelvic pain symptoms. VEGF-versus-saline differential translation was evaluated across all 12 TRAP-IP samples (N=3 per sex per group) and restricted to autosomal protein-coding genes at nominal p<0.05 with >1.5-fold change. These criteria identified 566 genes, including immune, lipid, ECM, GPCR/signaling, and transcription/translation related genes, with 327 upregulated and 239 downregulated in VEGF-instilled animals when compared to saline-instilled littermates (Fig. 5A, Supplemental Table S3). Vesicle and adaptor trafficking genes *(Ap4m1, Rab40b, Lamp5, Ap1s3, Cope*) and mitochondrial genes (*Timm23, Gpx4, Ndufs2, Mff, Tomm70a*) were more abundant in TRAP-IP isolated from mice instilled with VEGF, whereas cytokine and immune genes (*Ccl9, Tnfsf14, Cxcl3, Ccl11, Ifna4*) and additional immune-related genes (A*dgre1, Ccr4, Selplg, Tnfaip6, Irf1*) were more abundant in saline-instilled animal SGCs (Fig. 5C). Taken together, our data demonstrated a clear shift in SGC translatome in CPP conditions, from baseline functions (including the sex-specific ECM/lipid gene translation and immune function) to mitochondrial and vesicular gene translation.

**Figure 5.**
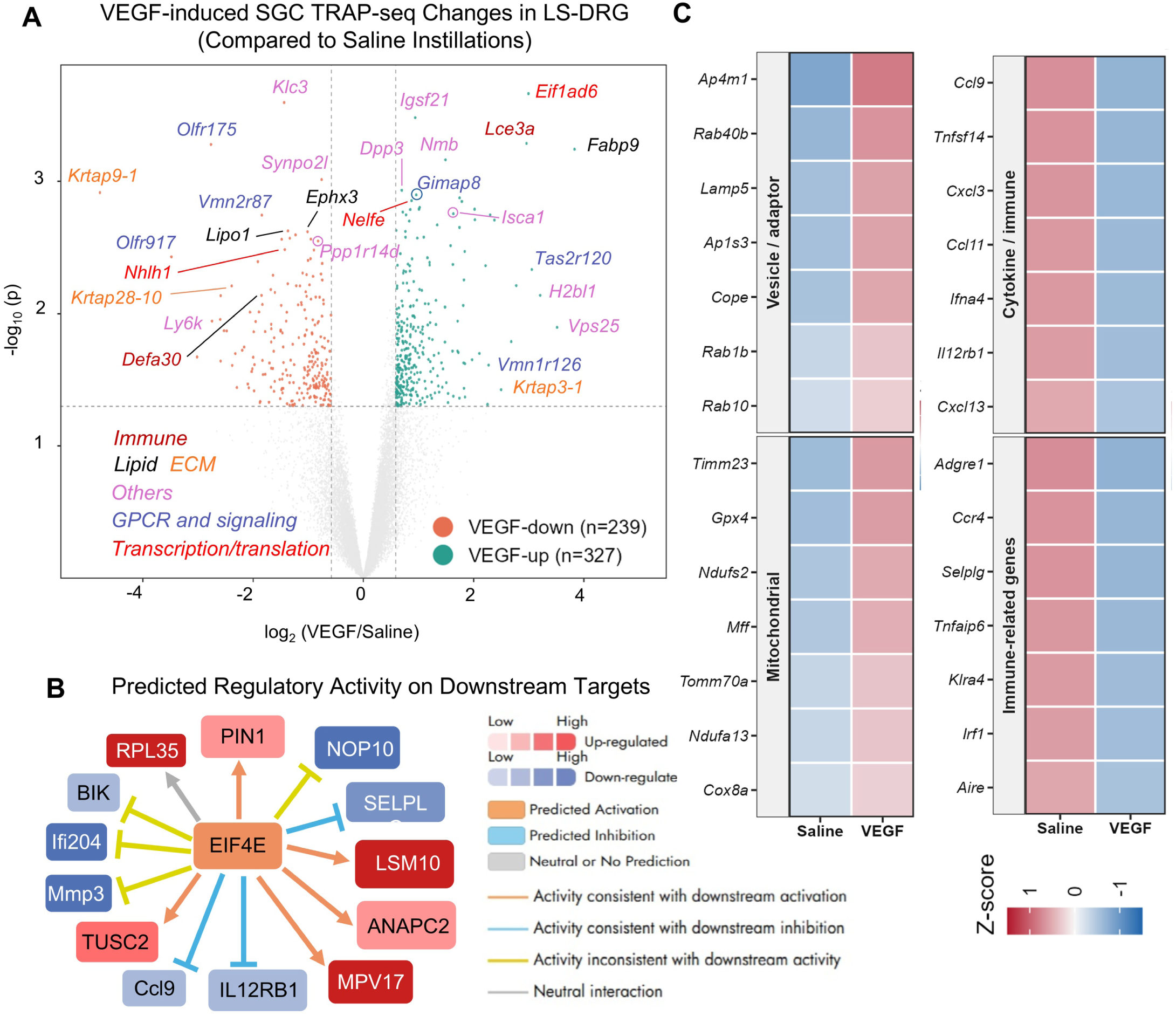
VEGF treatment reprograms the SGC translatome through an EIF4E-centered translational regulatory network. (A) Volcano plot of differential gene expression in the Sox10-TRAP-seq SGC translatome following VEGF versus saline treatment (protein-coding, autosomal genes; DESeq2, nominal *p*<0.05, >1.5-fold change in LS-DRG (LS-DRG). VEGF significantly downregulated 239 genes (orange) and upregulated 327 genes (green) relative to saline; Non-significant (grey). (B) Ingenuity Pathway Analysis (IPA) upstream regulator analysis identified EIF4E as the sole predicted activated translational regulator among differentially expressed genes Edge color indicates whether downstream target activity is consistent (orange) or inconsistent (yellow) with the predicted EIF4E activation state; node fill indicates observed up-(red) or down-regulation (blue). (C) Heatmaps of selected vesicle/adaptor, mitochondrial, cytokine/immune and immune-related genes, shown as row Z-scores of mean [mean of per-sample VST z-scores] in saline and VEGF groups

We next investigated what factors might contribute to the CPP-associated translational regulation. Ingenuity Pathway Analysis (IPA) of all 566 genes predicted *Eif4e* as an upstream translational regulator which was activated in TRAP-IP isolated from VEGF-induced animals. The downstream targets of EIF4E include *Rpl35, Lsm10, Anapc2, Mpv17, Tusc2* which were activated, and *Nop10, Selplg, Il12rb1, Ccl9* which were inhibited in VEGF-instilled animal SGCs (Fig. 5B). eIF4E binds the mRNA 5′ cap and is rate-limiting for cap-dependent translation. Together, these findings raise the possibility that EIF4E-dependent translational regulation in SGCs contributes to the development or maintenance of CPP.

### VEGF instillations induce limited cytokine changes in LS-DRG only in females

Under inflammatory and nerve-injury conditions, SGCs have been shown to release pro-inflammatory cytokines^42,43^. Due to the translatome pattern shift, we hypothesized that the cytokine level in LS-DRG may be decreased following VEGF instillations. To test this hypothesis, cytokines were measured with a 23-plex bead assay in whole LS-DRG lysates from a separate cohort of naïve, saline-instilled and VEGF-instilled C57BL/6J mice (n = 5–7 per sex per group; STAR Methods). 17 analytes were quantifiable, including pro-inflammatory cytokines (IL-1a, IL-6, IL-12 (p40), IL-12 (p70), IL-17A, TNF, and IFNγ, Fig. 6A), chemokines (KC, RANTES, MCP-1, MIP-1a, Fig. 6B), Type 2 immunity related cytokines (IL-4, IL-5, Eotaxin, Fig. 6C), regulatory cytokines (IL-10, Fig. 6D), and growth & survival factors (IL-3 and IL-2, Fig. 6E). Our data showed that 3 out of 17 analytes were differentially expressed in female LS-DRG. IL-17A was significantly lower in both saline-instilled and VEGF-instilled females than naive females (N=3) (p<0.01 and p<0.001, respectively). IFNγ was also lower in both instilled groups than in naive females (p<0.01 for both comparisons). IL-10 was lower in VEGF-instilled females than naïve females (p < 0.05). These changes were likely associated with instillation procedure rather than VEGF-induced bladder inflammation. The other 14 analytes, including MCP-1, MIP-1a, IL-2, and TNF were not differentially expressed among three groups. No sex differences were detected for any analytes across three experimental conditions.

**Figure 6.**
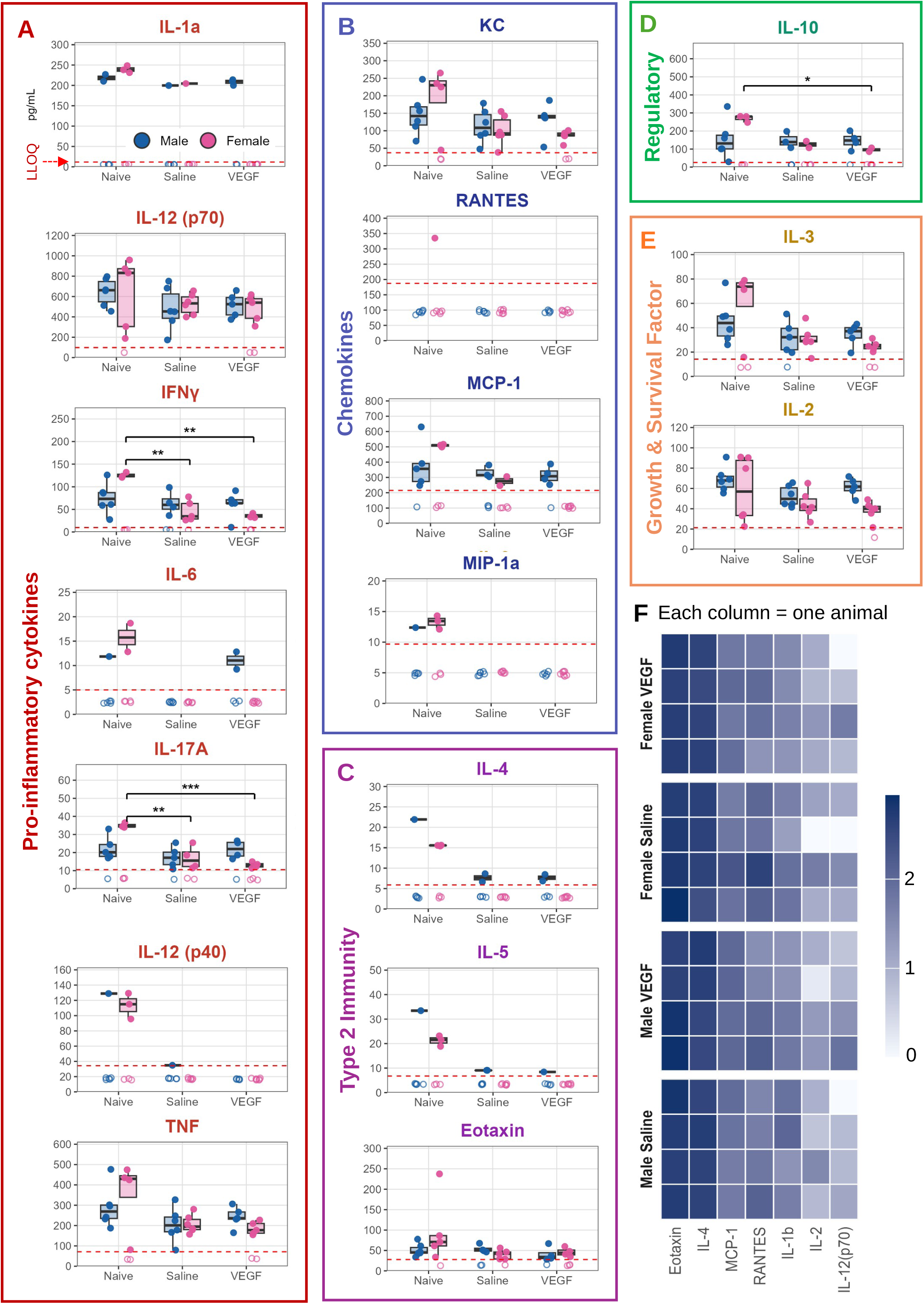
Cytokine and chemokine levels in LS-DRG. Bio-Plex Pro mouse 23-plex measurements in DRG homogenates from naïve, saline-instilled and VEGF-instilled male (blue) and female (pink) mice. Analytes are grouped as (A) pro-inflammatory cytokines, (B) chemokines, (C) type 2 immunity, (D) regulatory and (E) growth and survival factors. Each point is one animal. Filled symbols are above and open symbols below the lower limit of quantification (LLOQ, red dashed line). Boxes show the median and interquartile range. *p < 0.05, **p < 0.01, ***p < 0.001 (F) Transcript levels of Bio-Plex target genes in bulk LS-DRG RNA-seq from a separate, previously published cohort^6^ (n = 4 per sex × treatment), so protein and transcript data are not paired within animals. Shown are the 7 of 23 target genes meeting the detection criterion (mean normalized count ≥10 and ≥3 samples with raw count ≥5), plotted as log₁₀(normalized count + 1); each row is one animal. No gene differed between VEGF and saline (DESeq2, ∼sex + treatment, p ≥ 0.05).

To confirm these cytokines are transcribed in LS-DRG, we reanalyzed previously published bulk LS-DRG RNA-seq data^6^ (from the same experimental time point). Because the transcriptomic and protein datasets were generated from different cohorts, transcript and protein measurements are not paired within animals (Fig. 6F). Seven of the genes encoding the 23 Bio-Plex targets met the prespecified detection criterion (STAR Methods) and are shown for each animal as log10(normalized count+1); two of these, Il4 and Il2, are not supported by read-level inspection (see Discussion). None of the seven detected transcripts differed between VEGF- and saline-instilled animals (Fig. 6F).

### Genes related to canonical SGC-neuron interactions are not altered by VEGF

SGCs express pathways known to regulate neuronal excitability and activity, such as neurotransmitter uptake and recycling, ion homeostasis, and antigen presentation^9^. Here, we examined whether VEGF instillations induced changes in genes related to canonical SGC functions. Due to the limitation of ribosomal-associated mRNA recovered in TRAP-IP fractions, we first used the scRNA-seq and snRNA-seq datasets^14,30^ to identify key SGC functions related to neuronal excitability and network activity. SGC-specific genes were identified based on: 1) mean SGC cluster expression at or above the 33rd percentile of all genes, and 2) mean non-SGC cluster expression below 2.5-fold of the mean SGC cluster expression. This analysis resulted in a total of 133 genes in 15 functional categories (Supplemental Fig. S4), including K^+^ buffering, gap and tight junctions, glutamate/GABA transport and synthesis, ATP and adenosine signaling, NFκB signaling, cytokine production, and antigen presentation. We next compared the expression level of these 133 genes between TRAP-IP isolated from saline- and VEGF-instilled animals. Our data showed that all 133 genes remained unchanged (FDR>0.05 for either term; Supplementary Table S5). Specifically, *Gfap, Slc1a6,* and *Psmb8* met the study-wide threshold of nominal p<0.05 with >1.5-fold change for the treatment effect, while *Cldn12* and *Casp1* met this threshold for the sex × treatment interaction. Together, these data suggested that the translation level of genes related to canonical sensory SGC functions remained largely unchanged under CPP conditions at the time point examined in our study, although significant transcriptional or translational level changes may have occurred before this time point when the bladder was first exposed to VEGF.

### Sex-divergent hormone receptor signaling in the SGC translatome

We next asked whether sex hormone signaling drives translatome sex differences. We first determined if *Ar* (AR)*, Esr1* (ER1), and *Esr2* (ER2) are expressed in sensory SGCs using two independent DRG datasets snRNA (both sexes)^30^ and scRNA (only males)^14^, with SGC clusters pooled within platform and using neurons as reference and *Plp1* and *Fabp7* as SGC marker controls (Fig. 7A). *Ar* and *Esr1* were detected in only 0.3–3.4% of SGCs or SGC nuclei, compared with 3.6–81% of neurons, and *Esr2* was essentially absent from SGCs (0–0.09%). The SGC markers were also detected in most neuron-cluster cells, illustrating ambient RNA in droplet data. Because *Esr2* was essentially undetected, only *Esr1* and *Ar* were carried forward (Fig. 7B–D).

**Figure 7.**
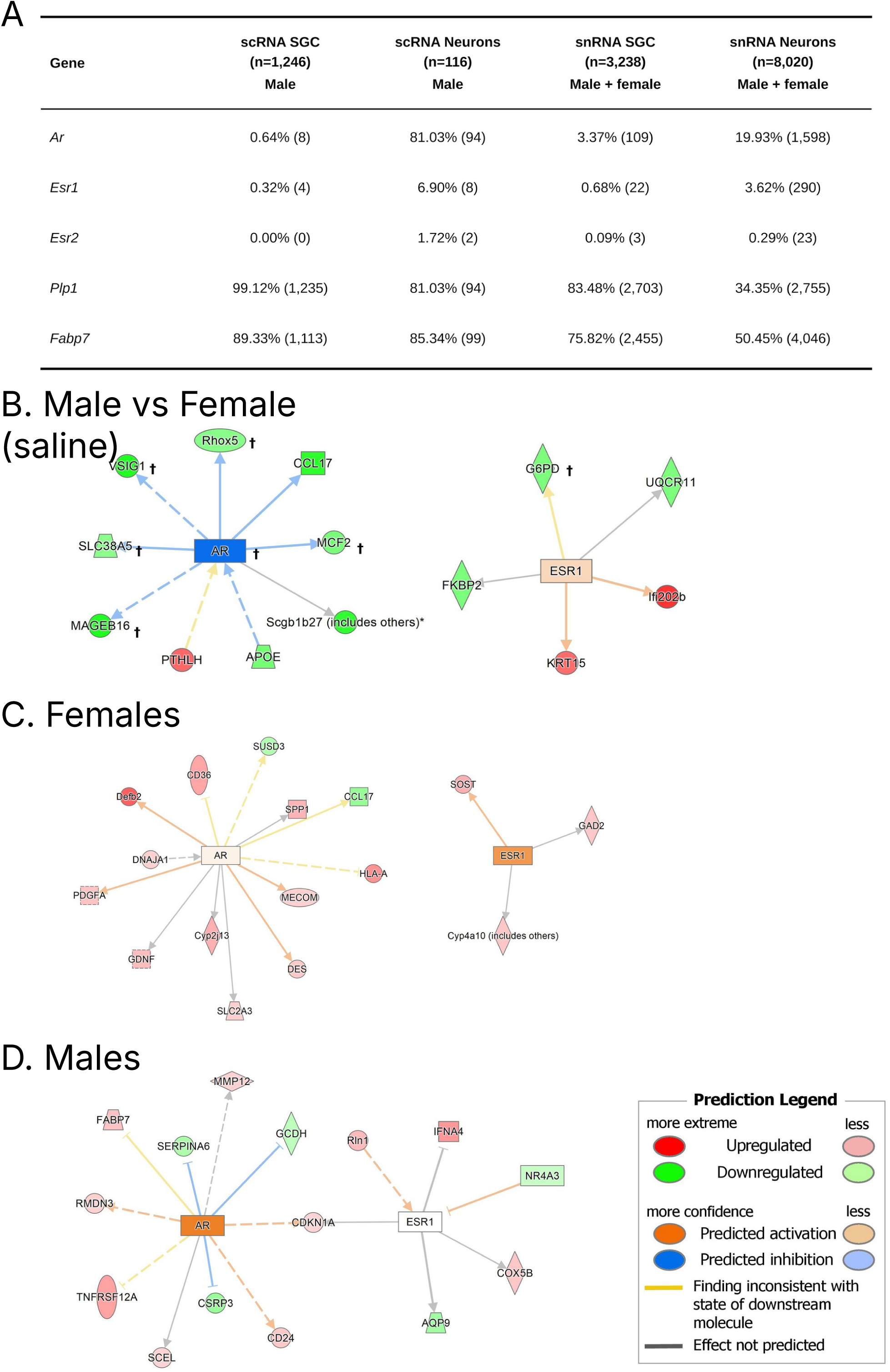
Sex hormone receptor detection and predicted receptor activity in the SGC translatome. (A) Detection of *Ar*, *Esr1* and *Esr2* in two independent DRG single-cell datasets, with satellite glial clusters pooled within platform and neurons as reference. Values are the percentage of cells or nuclei with non-zero counts (absolute counts in parentheses). *Plp1* and *Fabp7* are shown as satellite glial markers. No replicate-level data were available; the analysis is descriptive. (B) *Ar*/*Esr1*-centred curated network overlaid with saline-only male versus female translatome fold changes (n=3 per sex), showing baseline sex differences in the absence of VEGF. Node fill and edge style follow the Prediction Legend; expression colors indicate the direction of change in males relative to females. †X-linked (*Ar*, *Rhox5*, *Vsig1*, *Slc38a5*, *Mageb16*, *Mcf2*, *G6pd*); in a male–female comparison, female-biased expression of X-linked genes reflects X-chromosome dosage in addition to any receptor-mediated regulation (C, D). The same curated network overlaid with VEGF versus saline fold changes in females (C) and males (D), analyzed separately (p<0.05, n=3 per group). Neither receptor was itself differentially translated in either sex. Based on the direction of downstream target changes, IPA predicted ESR1 activation in females and AR activation in males.

To determine whether *Esr1* or *Ar* signaling drives sex-specific SGC translatome pattern at the baseline, TRAP-IP from saline-instilled animals were compared between sexes and overlaid on an *Ar* and *Esr1*-centred curated network using IPA (Fig. 7B). Among all 16 differentially expressed genes, 7 are X-linked (*Ar*, *Rhox5*, *Vsig1*, *Slc38a5*, *Mageb16*, *Mcf2*, *G6pd*) and showed significant female-biased expression. Based on the direction of downstream target changes, IPA predicted *Ar* inhibition and *Esr1* activation in male relative to female SGC translatome.

Lastly, we asked if *Esr1* and *Ar* signaling contributes to VEGF-induced translatome changes in male and female mice. Female and male gene lists (p<0.05) were analyzed separately in IPA with *Esr1* or *Ar* as reference nodes (Fig. 7C-D). Neither receptor was differentially translated in VEGF-instilled TRAP-IP when compared to saline TRAP-IP. However, based on downstream gene expression changes, IPA predicted *Esr1* signaling activation in female SGC translatome and *Ar* signaling activation in male SGC translatome. These pathway-based predictions are derived from downstream target changes rather than direct measurements of receptor activity and therefore do not necessarily indicate hormonal causality.

## Discussion

This study used TRAP-seq to profile *Sox10+* SGC translatome in LS-DRG from mice with VEGF-induced pelvic hypersensitivity. Three principal findings emerged from our study: 1) the immunoprecipitated fraction was enriched for canonical satellite glial transcripts and depleted of neuronal, fibroblast, endothelial, pericyte, and immune markers. 2) the baseline satellite glial translatome was sex dependent, yet 98.1% of these sex differences were not detected in paired whole-DRG RNA (Input). 3) VEGF shifted the translatome toward vesicular and mitochondrial programs and away from cytokine- and immune-related programs, with EIF4E predicted as an activated upstream regulator. These translational changes occurred without evidence of broad inflammatory activation across the 23 cytokines and chemokines measured in whole-DRG lysates.

### Sox10-TRAP for SGC translatome study

Sox10 is required for the development of both SGCs and Schwann cells^44^, so a Sox10-driven ribosomal tag should label both populations. In the *Sox10 CreER^T2^::L10a-GFP* mice, tamoxifen-induced recombination labeled largely SGCs in neuronal soma-rich region. Marker-panel analysis using two independent DRG single-cell and single-nucleus atlases showed enrichment of SGC markers in the TRAP-IP and depletion of all six non-SGC marker classes (Fig. 1). To our knowledge, this is the first study to define satellite glial translatome. The key advantage here is direct tissue homogenization, which avoids cell dissociation and makes the morphology of SGCs less limiting for molecular profiling.

Besides avoiding cell dissociation, we also found that TRAP-seq is very effective in detecting low-expressing transcripts. We examined gap-junction and GPCR-signaling genes that are known to participate in SGC functions when compared to other sequencing methods (Fig. 2). Several signaling transcripts readily detected in the TRAP-IP were low or undetected in whole-DRG input samples and published naïve SGC pseudobulk datasets^30^. This is not simply because SGC-restricted transcripts were diluted in the Input, as the pseudobulk analysis was restricted to SGC nuclei. We think that the low-copy transcripts are particularly vulnerable to incomplete capture in snRNA-seq^19,45^ due to sample preparation, cell recovery, and normalization^30,45^.

### GPCRs in SGC translatome suggest SGC function

Our data showed that Sox10-TRAP recovers SGC transcripts. However, abundance in the TRAP-IP alone does not establish SGC origin. We therefore began with all curated GPCRs in the IUPHAR/BPS Guide to Pharmacology^46^ with three independent lines of evidence: TRAP-IP abundance above an empirical floor defined by presumed negative-control chemosensory receptor genes, consistent expression in published bulk RNA-seq of FACS-sorted naïve SGCs^13^, and no clear attribution to other cell types in two DRG atlases. Thirty-two of 343 receptors met all three criteria (Fig. 3A), which is described as SGC-expressed (rather than SGC-specific). Nineteen of the 32 receptors belong to the adhesion (9), Frizzled/Smoothened (6), Celsr (2), and LGR (2) families, which are associated with cell contact, tissue polarity, and developmental signaling. Adhesion GPCRs bind cell-surface and extracellular-matrix ligands^47^; The expression of these GPCRs in SGC translatome this composition suggested sensing or maintenance of the glia–neuron interface. The repertoire also includes receptors for GABA (*Gabbr1*), adenosine (*Adora1*), endothelins, PACAP (*Adcyap1r1*), and lysophospholipids. *Ednrb* and *Gpr37l1* are among the most abundant GPCR transcripts in sorted SGCs, matching their reported high expression in mouse and human DRG. Specifically, GPR37L1 has been localized to SGCs and linked to potassium channel regulation and resolution of neuropathic pain^48^. *Gpr37l1*, the shortest receptor in the repertoire (2.3 kb, about one-third of the median), lay closest to the detection floor (Fig. 3B) despite its high expression in sorted SGCs and mouse DRG, consistent with the length bias of CPM^49^. The TRAP-IP-to-Input ratio (Fig. 3C) describes relative RNA representation in the TRAP and input fractions, not cell specificity or receptor activation. The repertoire was stable when VEGF libraries were excluded (Supplementary Fig. S3). *Lhcgr* was an unexpected member: it met all three criteria and was detected in all four sorted-glia replicates, although luteinizing hormone signaling in SGCs has not been examined.

### Male and female SGCs signaling differently at the baseline but not in CPP condition

In pain-free animals (saline-instilled), 2,229 genes were differentially expressed between male and female mice in TRAP-IP, but only 43 of them were differentially expressed in the Input between sexes (Fig. 4). This data suggested that male and female SGCs exhibit profound sex difference in baseline signaling events, possibly more pronounced than those observed in the whole DRG. On the other hand, the canonical genes in SGC translatome were largely indifferent between sexes (40 of 43 genes, 93%) either at the baseline or following VEGF instillations, suggesting that most core SGC marker genes were not strongly sex-associated.

We next focused our functional interpretation on autosomal protein-coding genes in pathway analysis. Noncoding transcripts can associate with ribosomes without encoding proteins, and their recovery by TRAP therefore does not indicate assay failure^50^. X- and Y-linked genes were excluded because their differential expression may reflect sex-chromosome complement or dosage rather than autosomal SGC programs^51^. Female-biased genes were enriched for extracellular-matrix and remodeling programs, whereas male-biased genes were enriched for lipid and eicosanoid signaling. Sex-divergent immune mechanisms have been established in DRG using mouse pain models^7^ including our own CPP mouse models^6^. However, sex differences in this study were largely restricted to baseline conditions but not in mice exhibiting VEGF-induced pelvic pain symptoms (Fig. 4).

### Potential activation of EIF4E, a translational regulator in CPP condition

IPA predicted EIF4E activation based on a network of 13 differentially abundant transcripts (Fig. 5B): increased *Anapc2, Lsm10, Mpv17, Pin1, Rpl35,* and *Tusc2*, and decreased *Bik, Ccl9, Ifi206, Il12rb1, Mmp3, Nop10,* and *Selplg*. MNK1/2–eIF4E axis contributes to injury-induced plasticity and chronic pain^25^, and the selective MNK1/2 inhibitor tomivosertib reverses neuropathic pain in mice^52^. EIF4E phosphorylation regulates translation of a defined subset of DRG mRNAs including *Bdnf*^26^, and MNK1–eIF4E signaling can promote mTORC1 activity through translational regulation of RagA^23^. Although these studies used sensory neurons rather than Sox10-positive glia, the present findings raise the possibility that VEGF alters translational regulation in Sox10-positive DRG cells. However, *Eif4e* gene translation was not changed between VEGF and saline conditions. Establishing an EIF4E-dependent mechanism will require direct measurement of eIF4E abundance, phosphorylation, or downstream translational activity.

### The SGC translatome shift towards mitochondrial and vesicular functions in CPP

Figure 5C summarizes VEGF-induced transcripts changes in SGC translatome. Transcripts in vesicle/adaptor-trafficking, mitochondrial, cytokine/immune, and other immune-related groups were differentially expressed. The trafficking and mitochondrial transcripts shown were increased in SGCs from VEGF-instilled animals, while the immune-related transcripts were significantly decreased. These groupings illustrate individual differences rather than coordinated pathway regulation. Rotation gene set testing^53^ found no significant VEGF-associated changes in the prespecified trafficking or mitochondrial gene sets. Although 17 of the 21 gene sets tested had positive mean shifts, this shared direction is attributable to a global upward offset. The reduction in immune-associated transcripts nevertheless prompted us to ask whether the cytokines measured at the protein level showed corresponding whole-DRG transcript patterns.

### SGC translatome in CPP conditions shows immune suppression, while cytokine analysis in whole DRG does not reflect VEGF-induced changes

Because immune-related transcripts decreased in the SGC translatome, we asked whether the DRG showed corresponding cytokine changes. Seventeen of 23 analytes were quantifiable above their assay-specific LLOQ in at least one group, and values below the LLOQ should not be read as absence of protein. All significant differences were between naïve and instilled females (lower IFN-γ and IL-17A after both instillations; lower IL-10 after VEGF only), with none between saline and VEGF, so they most likely reflect the instillation procedure rather than VEGF. In independent bulk RNA-seq from the same model^6^, read-level inspection showed that the high Il4 and Il2 counts arose from non-exonic reads and an internal-priming stack, respectively, leaving five reliably transcribed analytes, none of which changed with VEGF. Because protein and transcript data come from different animals and whole-DRG tissue, they provide independent context rather than paired validation and cannot identify the cellular source. Together, they show no broad VEGF-associated inflammatory response in the DRG, consistent with the female-specific immune-resolution signature in our earlier bulk analysis^6^. VEGF-associated recruitment of myeloid-derived suppressor cells, reported in ovarian cancer, is one mechanism that could be tested. VEGF-associated recruitment of myeloid-derived suppressor cells was reported in ovarian cancer^54^, presenting a potential mechanism for future testing.

### Sex hormonal signaling does not directly contribute to the baseline sex differences in SGC translatome

Our data showed that baseline SGC translatome was sex-dependent (Fig. 4). Therefore, we asked whether *Ar* and *Esr1* known targets changed coherently using receptor-seeded IPA networks (nominal P<0.05, N=3 per group; no gene passed FDR<0.05)^55^. At baseline, lower predicted AR activity in males depended mainly on X-linked targets (5 of 6; Fig. 7B) and so likely reflects X-chromosome dosage rather than androgen signaling. Following VEGF treatment, only the male AR network leaned toward activation (6 consistent and 2 opposing changes; z score<2, below threshold; Fig. 7D). The female AR network was evenly split, and neither ESR1 network yielded a prediction (Fig. 7C). Thus, no coherent receptor-driven response was detected in either sex. *Esr2* was not examined because it was rarely detected (0-0.1%) in a male single-cell atlas and a combined-sex single-nucleus atlas^30^ (Fig. 7A). In addition, a published bulk RNA-seq of FACS-sorted SGC data showed that *Ar* and *Esr1* were below 1 FPKM in all four replicates, and *Esr2* were near zero.

Cancer studies have linked sex hormone receptor signaling to translational regulation. In prostate cancer models, AR increases 4E-BP1, thereby limits eIF4F complex assembly.^56^. In ER-positive breast cancer models, eIF4E can promote estrogen independence^57^ and increase ERα and FOXM1 translation^58^. Whether similar interactions contribute to sex differences in the SGC translatome remains unknown, and the low *Ar* and *Esr1* expression in SGCs limited support for a direct receptor-mediated mechanism.

### Study limitations

This study has several limitations. First, *Sox10* drives ribosomal tag expression in some Schwann cells. Second, immunoprecipitates from L10-GFP-negative mice was not performed, so non-specific RNA binding could not be quantified or subtracted. Third, all measurements were performed on day 16 after the first instillation, which does not capture early glial response. Finally, the translatome findings are transcript-level observations. Future study is needed to measure protein abundance, localization, and activity.

## Supporting information

Supplemental Table S1 - Sex differences in the SGC translatome

Supplemental Table S2 - Autosomal protein-coding sex-biased genes

Supplemental Table S3 - VEGF-associated translatome differences

Supplemental Table S4 - DRG cytokine and chemokine measurements

Supplemental Table S5 - SGC functional gene panel

---

**Supplemental Table S1. Sex differences in the SGC-enriched translatome of saline-instilled mice, related to Figure 4**. DESeq2 results (∼batch + sex; n = 3 per sex) for 2,229 genes meeting nominal p < 0.05 and |log₂FC| > log₂(1.5): 1,255 male-biased and 974 female-biased. Positive log₂FC indicates higher male abundance. Genes comprise 768 autosomal protein-coding, 1,179 noncoding and 282 X/Y-linked genes. All genes are enriched in IP versus paired input.

**Supplemental Table S2. Autosomal protein-coding genes with sex differences in saline-instilled mice, related to Figure 4C**. Gene list containing 768 autosomal protein-coding genes: 527 male-biased and 241 female-biased. Twenty-three meet BH-adjusted p < 0.05. Noncoding and X/Y-linked genes were excluded from this functional interpretation subset.

**Supplemental Table S3. VEGF-associated differences in the SGC-enriched translatome, related to Figure 5**. DESeq2 results (∼batch + sex + treatment; n = 6 per treatment, 3 per sex) for 566 autosomal protein-coding genes meeting nominal p < 0.05 and greater than 1.5-fold change in either direction at day 16. Positive log₂FC indicates higher abundance with VEGF (327 UP; 239 DOWN). This exploratory list was used for IPA upstream regulator analysis.

**Supplemental Table S4. DRG cytokine and chemokine measurements, related to Figure 6**. Bio-Plex Pro mouse 23-plex measurements in LS-DRG homogenates from naïve, saline-instilled and VEGF-instilled male and female mice, assayed in duplicate. MV4 (male VEGF) was excluded because fluorescence for all 23 analytes was at or near blank-well background, indicating a failed sample (final n = 6 per group; male VEGF n = 5, female VEGF n = 7).

**Supplemental Table S5. VEGF-associated and sex × treatment effects in the SGC functional gene panel, related to Supplemental Fig. S4.** The table gives counts per functional category for 133 SGC-expressed genes (138 category assignments). Effects were estimated by DESeq2 (∼batch + sex + treatment, plus a sex:treatment term for the interaction; n = 3 per sex per treatment). The p < 0.05 & >1.5-fold columns report nominal thresholds. FDR is Benjamini–Hochberg corrected across the 133 panel genes.

## STAR Methods

### KEY RESOURCES TABLE

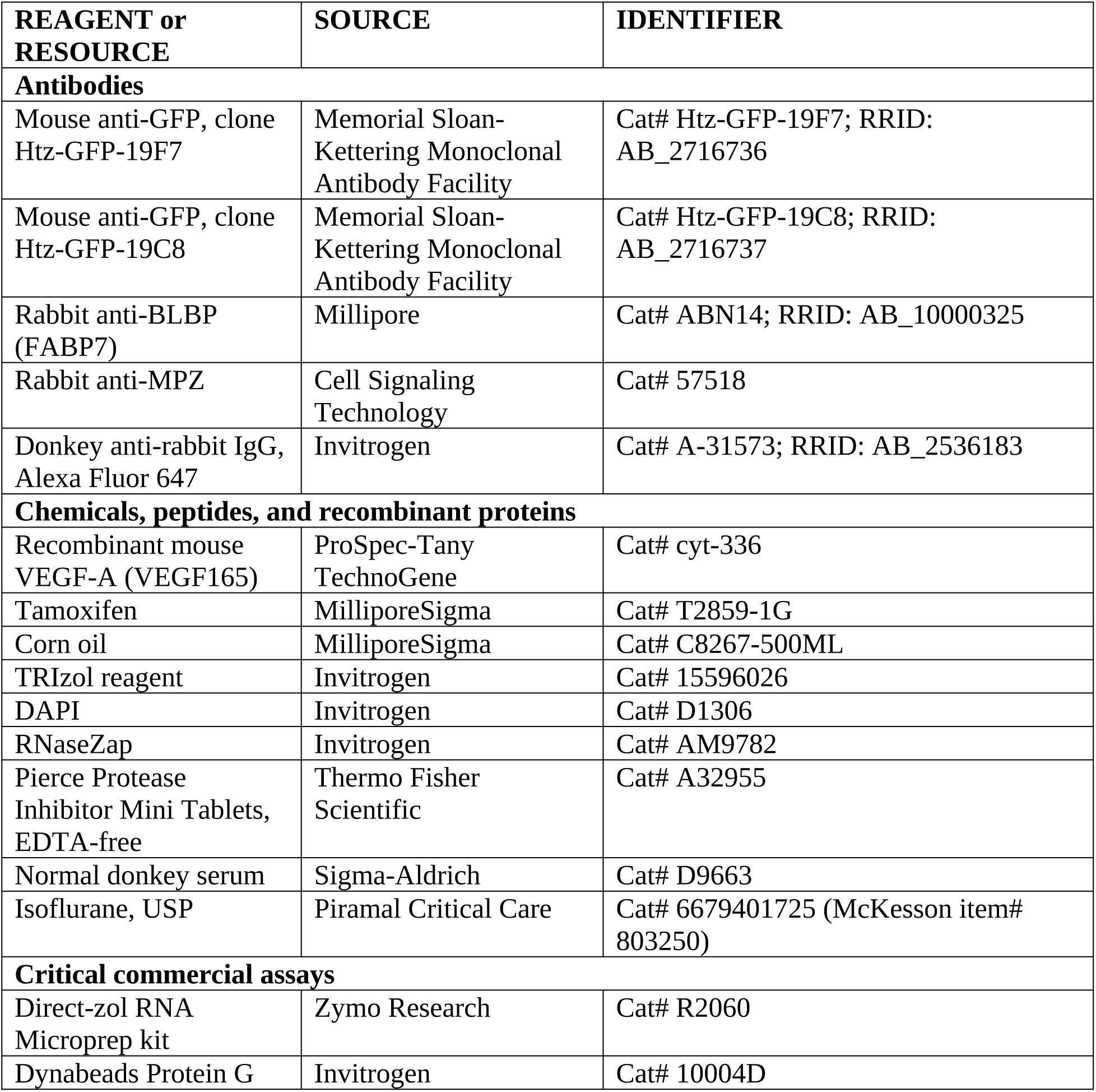
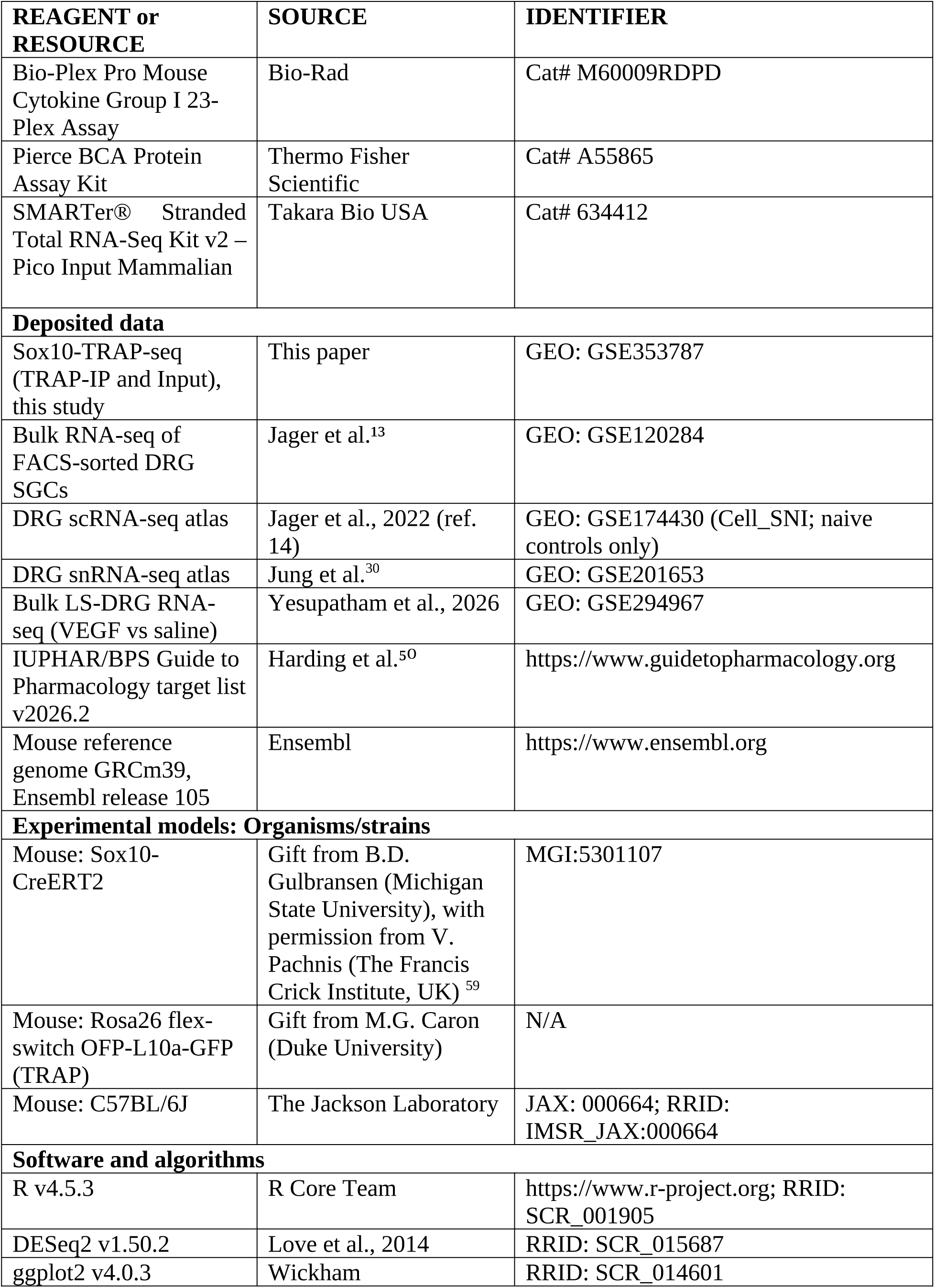
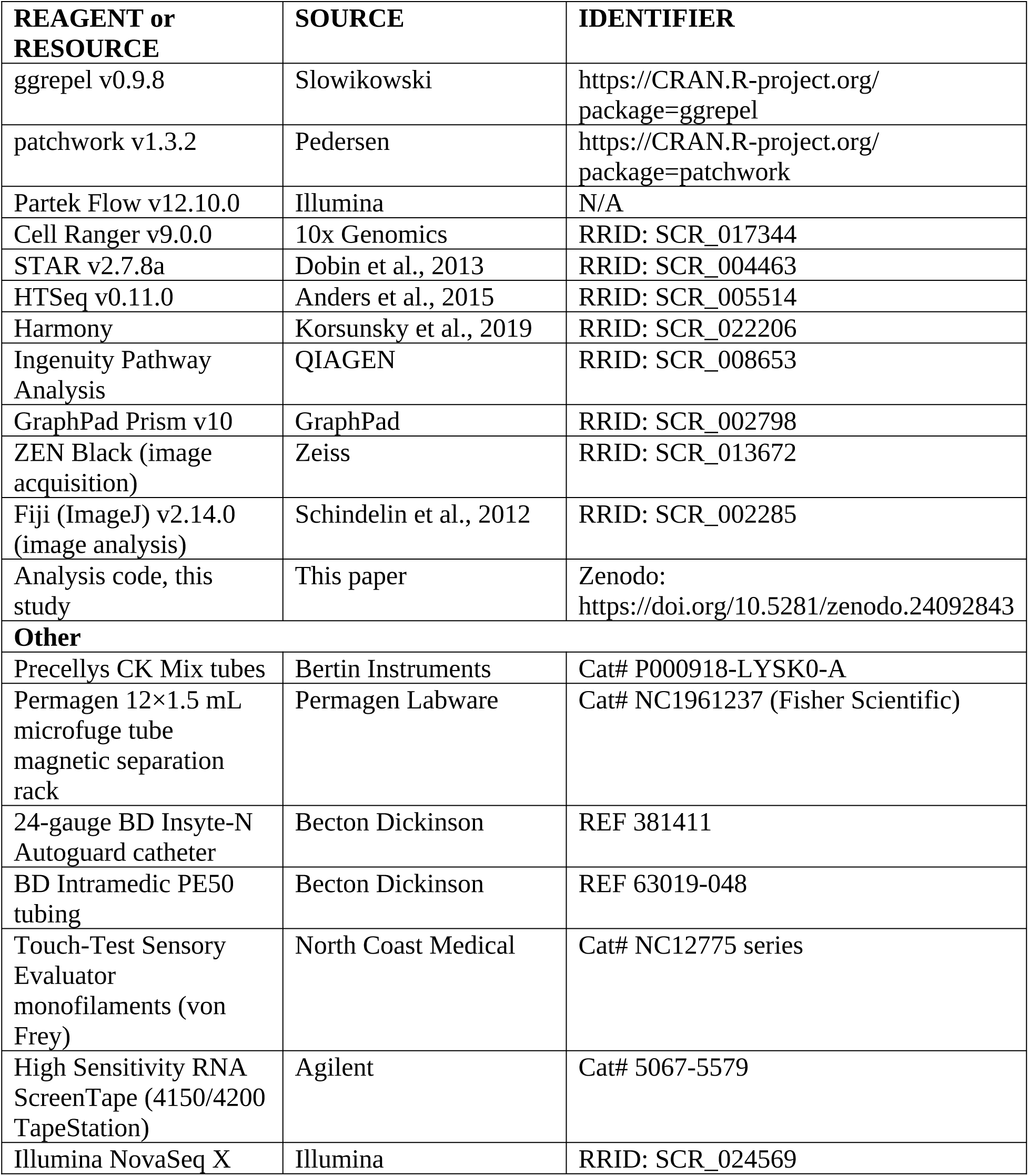

| **REAGENT or RESOURCE** | **SOURCE** | **IDENTIFIER** |
|---|---|---|
| **Antibodies** |  |  |
| Mouse anti-GFP, clone Htz-GFP-19F7 | Memorial Sloan-Kettering Monoclonal Antibody Facility | Cat# Htz-GFP-19F7; RRID: AB_2716736 |
| Mouse anti-GFP, clone Htz-GFP-19C8 | Memorial Sloan-Kettering Monoclonal Antibody Facility | Cat# Htz-GFP-19C8; RRID: AB_2716737 |
| Rabbit anti-BLBP (FABP7) | Millipore | Cat# ABN14; RRID: AB_10000325 |
| Rabbit anti-MPZ | Cell Signaling Technology | Cat# 57518 |
| Donkey anti-rabbit IgG, Alexa Fluor 647 | Invitrogen | Cat# A-31573; RRID: AB_2536183 |
| **Chemicals, peptides, and recombinant proteins** |  |  |
| Recombinant mouse VEGF-A (VEGF165) | ProSpec-Tany TechnoGene | Cat# cyt-336 |
| Tamoxifen | MilliporeSigma | Cat# T2859-1G |
| Corn oil | MilliporeSigma | Cat# C8267-500ML |
| TRIzol reagent | Invitrogen | Cat# 15596026 |
| DAPI | Invitrogen | Cat# D1306 |
| RNaseZap | Invitrogen | Cat# AM9782 |
| Pierce Protease Inhibitor Mini Tablets, EDTA-free | Thermo Fisher Scientific | Cat# A32955 |
| Normal donkey serum | Sigma-Aldrich | Cat# D9663 |
| Isoflurane, USP | Piramal Critical Care | Cat# 6679401725 (McKesson item# 803250) |
| **Critical commercial assays** |  |  |
| Direct-zol RNA Microprep kit | Zymo Research | Cat# R2060 |
| Dynabeads Protein G | Invitrogen | Cat# 10004D |
| **REAGENT or RESOURCE** | **SOURCE** | **IDENTIFIER** |
| Bio-Plex Pro Mouse Cytokine Group I 23-Plex Assay | Bio-Rad | Cat# M60009RDPD |
| Pierce BCA Protein Assay Kit | Thermo Fisher Scientific | Cat# A55865 |
| SMARTer® Stranded Total RNA-Seq Kit v2 – Pico Input Mammalian | Takara Bio USA | Cat# 634412 |
| **Deposited data** |  |  |
| Sox10-TRAP-seq (TRAP-IP and Input), this study | This paper | GEO: GSE353787 |
| Bulk RNA-seq of FACS-sorted DRG SGCs | Jager et al.<sup>13</sup> | GEO: GSE120284 |
| DRG scRNA-seq atlas | Jager et al., 2022 (ref. 14) | GEO: GSE174430 (Cell_SNI; naive controls only) |
| DRG snRNA-seq atlas | Jung et al.<sup>30</sup> | GEO: GSE201653 |
| Bulk LS-DRG RNA-seq (VEGF vs saline) | Yesupatham et al., 2026 | GEO: GSE294967 |
| IUPHAR/BPS Guide to Pharmacology target list v2026.2 | Harding et al.<sup>50</sup> | https://www.guidetopharmacology.org |
| Mouse reference genome GRCm39, Ensembl release 105 | Ensembl | https://www.ensembl.org |
| **Experimental models: Organisms/strains** |  |  |
| Mouse: Sox10-CreERT2 | Gift from B.D. Gulbransen (Michigan State University), with permission from V. Pachnis (The Francis Crick Institute, UK)<sup>59</sup> | MGI:5301107 |
| Mouse: Rosa26 flex-switch OFP-L10a-GFP (TRAP) | Gift from M.G. Caron (Duke University) | N/A |
| Mouse: C57BL/6J | The Jackson Laboratory | JAX: 000664; RRID: IMSR_JAX:000664 |
| **Software and algorithms** |  |  |
| R v4.5.3 | R Core Team | https://www.r-project.org; RRID: SCR_001905 |
| DESeq2 v1.50.2 | Love et al., 2014 | RRID: SCR_015687 |
| ggplot2 v4.0.3 | Wickham | RRID: SCR_014601 |
| **REAGENT or RESOURCE** | **SOURCE** | **IDENTIFIER** |
| ggrepel v0.9.8 | Slowikowski | https://CRAN.R-project.org/package=ggrepel |
| patchwork v1.3.2 | Pedersen | https://CRAN.R-project.org/package=patchwork |
| Partek Flow v12.10.0 | Illumina | N/A |
| Cell Ranger v9.0.0 | 10x Genomics | RRID: SCR_017344 |
| STAR v2.7.8a | Dobin et al., 2013 | RRID: SCR_004463 |
| HTSeq v0.11.0 | Anders et al., 2015 | RRID: SCR_005514 |
| Harmony | Korsunsky et al., 2019 | RRID: SCR_022206 |
| Ingenuity Pathway Analysis | QIAGEN | RRID: SCR_008653 |
| GraphPad Prism v10 | GraphPad | RRID: SCR_002798 |
| ZEN Black (image acquisition) | Zeiss | RRID: SCR_013672 |
| Fiji (ImageJ) v2.14.0 (image analysis) | Schindelin et al., 2012 | RRID: SCR_002285 |
| Analysis code, this study | This paper | Zenodo: https://doi.org/10.5281/zenodo.24092843 |
| **Other** |  |  |
| Precellys CK Mix tubes | Bertin Instruments | Cat# P000918-LYSK0-A |
| Permagen 12×1.5 mL microfuge tube magnetic separation rack | Permagen Labware | Cat# NC1961237 (Fisher Scientific) |
| 24-gauge BD Insyte-N Autoguard catheter | Becton Dickinson | REF 381411 |
| BD Intramedic PE50 tubing | Becton Dickinson | REF 63019-048 |
| Touch-Test Sensory Evaluator monofilaments (von Frey) | North Coast Medical | Cat# NC12775 series |
| High Sensitivity RNA ScreenTape (4150/4200 TapeStation) | Agilent | Cat# 5067-5579 |
| Illumina NovaSeq X | Illumina | RRID: SCR_024569 |

### RESOURCE AVAILABILITY

#### Lead contact

Further information and requests for resources and reagents should be directed to and will be fulfilled by the lead contact, Sathish Kumar Yesupatham.

#### Materials availability

This study did not generate new unique reagents.

#### Data and code availability

TRAP-seq data (TRAP-IP and Input) have been deposited at GEO under accession GSE353787 and are publicly available as of the date of publication. This paper also analyzes existing, publicly available data: bulk RNA-seq of FACS-sorted DRG SGCs (GEO: GSE120284), DRG single-cell and single-nucleus RNA-seq atlases (GEO: GSE174430 and GSE201653, respectively), and bulk LS-DRG RNA-seq (GEO: GSE294967). Accession numbers are listed in the key resources table.

All original code has been deposited at Zenodo (https://doi.org/10.5281/zenodo.24092843) and is publicly available as of the date of publication.

Any additional information required to reanalyze the data reported in this paper is available from the lead contact upon request.

### EXPERIMENTAL MODEL AND STUDY PARTICIPANT DETAILS

#### Animals

To isolate ribosome-associated mRNA in the SGCs, mice harboring a Cre-inducible TRAP transgene (Rosa26 flex switch OFP-L10a-GFP) were crossed with the Sox10-CreERT2 mice^59^ for Sox10+ cell-specific expression of the TRAP transgene. Double transgenic mice were maintained as hemizygous for both Cre and the floxed allele (Sox10-CreERT2+/−::L10a-GFP+/f) and referred as “TRAP mice” hereinafter. The Sox10-CreERT2 line was a generous gift from Dr. Brian D. Gulbransen (University of Nevada, Reno), with permission from Dr. Vassilis Pachnis (The Francis Crick Institute, UK), who developed the line. Their Cre-littermate control (L10a-GFP+/f) mice were used as controls in some experiments. Mice were housed in a temperature- and humidity-controlled certified vivarium on a 14-hour/10-hour light/dark cycle, with ad libitum access to food and water. All experiments took place during the light cycle. All procedures were performed according to IACUC protocol 01147 approved by the IACUC of University of Colorado, Anschutz Medical Campus.

Both male and female mice were used. Mice were 8–12 weeks old at the start of experiments. Sex was included as a biological variable in the sex-stratified and combined analyses described below.

### METHOD DETAILS

#### Intravesical instillations in anesthetized animals

Intravesical instillations of mouse recombinant VEGF, VEGFA (also known as VEGF165) were performed to induce transient bladder inflammation and subsequent pelvic hypersensitivity, as previously described (Xie et al., 2022; Yesupatham et al., 2026). VEGFA was purchased from ProSpec-Tany TechnoGene Ltd. (Rehovot, Israel; catalog #cyt-336) and diluted in sterile saline (6.41 nM in 100 μL). Mice were anesthetized with isoflurane (1.5%) and transurethrally catheterized. Female mice were catheterized with sterile 24-gauge BD Insyte-N Autoguard Vialon catheter (Becton Dickinson, Sandy, Utah; REF 381411). Male mice were catheterized with sterile BD INTRAMEDIC Polyethylene Tubing/PE50 (Becton Dickinson, Sandy, Utah; REF 63019-048). All animals received three 100 µL instillations of VEGFA or saline during the two-week induction period. Intravesical instillations were performed via urethral catheter with a syringe attached to one end. To ensure consistent contact of the substances with the bladder lumen, and to avoid reflux or leakage, catheters were occluded and left in place for 30 minutes while the animals were kept on a warming pad under anesthesia. After 30 minutes, the catheter was removed and mice were allowed to recover from the isoflurane anesthesia and then returned to their home cage. Von Frey tests (VFT) were performed to assess visceral mechanical sensitivity before (day 0) and after the two-week induction period (day 15) on the same animals. DRG extractions were generally performed on Day 16.

#### Tamoxifen injections and DRG isolation

All experimental animals received intraperitoneal tamoxifen (40mg/ml in corn oil, 200mg/kg/day) on days 9 and 10 to induce L10a-GFP expression in SGCs. To minimize toxicity, the daily dose was divided into two injections of 100 mg/kg administered 12 hours apart. This is particularly important for male animals as injecting 200 mg/kg all at once can be lethal. On day 16, mice were sacrificed with isoflurane overdose and decapitated. LS-DRG (L1,L2,L6,S1,S2) were collected and rapidly placed in ice-cold dissection buffer and stored at −80°C for TRAP-seq experiments. For immunohistochemistry experiments, mice were perfused transcardiacally on day 16 and their LS-DRG were collected and processed as described in “Immunohistochemistry“.

#### Von Frey testing (VFT)

VFT were performed before instillations (day 0) and after (day 15). Mice were individually placed in clear plexiglass chambers (7.5 cm x 7.5 cm x 15 cm) equipped with wire grid floors (0.5 cm2 grid) and allowed to habituate until the extinction of exploratory behavior (1∼1.5 hours). A series of force-generating filaments (0.04, 0.16, 0.4, 1, 1.4, and 2g; North Coast Medical, Gilroy, California) were applied to the lower abdomen of the mice in the vicinity of the urinary bladder. Retraction of the abdomen, licking or scratching the stimulated area, or jumping were considered positive responses to the stimuli. Ten (10) repetitions were conducted with each filament and the percentage of positive responses to each force was calculated as withdrawal frequency (%).

#### Immunoprecipitation

TRAP Immunoprecipitation (TRAP-IP) was performed as previously described^24^ with slight modifications. All procedures were performed under strict RNase-free conditions; instruments and bench surfaces were cleaned with 70% ethanol and RNaseZap prior to use. For affinity matrix preparation, beads were prepared one day before the TRAP experiment. Dynabeads Protein G (200μL; Invitrogen, 10004D) were washed five times with 1× PBS containing 3% (w/v) IgG-free, protease-free BSA (Jackson ImmunoResearch, NC9871802), then washed three times in 0.15 M KCl IP wash buffer and incubated with 100μg total anti-GFP antibody (50μg clone Htz-GFP-19F7 and 50μg clone Htz-GFP-19C8; Memorial Sloan-Kettering Monoclonal Antibody Facility) in 200μL wash buffer for 2hours at room temperature with end-over-end rotation. Beads were washed and resuspended in 200μL wash buffer and stored at 4°C until use. The next day, DRGs were thawed on ice and transferred to Precellys CK Mix tubes (Cat. No.: P000918-LYSK0-A, Bertin Instruments) containing lysis buffer and homogenized using a Precellys Minilys homogenizer at high speed in 10-second intervals for a total of eight cycles in the cold room. A post-mitochondrial supernatant was obtained by centrifugation at 15,000 × g for 10 minutes at 4°C. An aliquot of 200 μL was reserved as the matched total RNA fraction (Input). The remaining fraction was added to the pre-prepared affinity matrix and incubated overnight at 4°C with end-over-end rotation. Beads were collected using a Permagen magnet. The unbound fraction was removed, and beads were washed four times with 0.35 M KCl IP wash. After the final wash, beads were resuspended in 100 µL TRIzol reagent (Invitrogen, 15596026) and incubated at 37 °C for 5 min with intermittent mixing to release ribosome-bound RNA. Beads were then separated on the magnet and the TRIzol lysate transferred to an RNase-free tube for RNA extraction.

#### RNA extraction, library preparation and sequencing

RNA was extracted from the TRAP-IP TRIzol eluate (100 µL, above) and from the Input fraction (1:3 sample:TRIzol) using the Direct-zol RNA Microprep kit (Zymo Research, R2060) according to the manufacturer’s instructions. RNA was eluted in 15 μL UltraPure DNase/RNase-free water. RNA was stored at −80°C until use. TRAP-IP yields were below 1 ng of total RNA per sample (High Sensitivity RNA ScreenTape, Agilent). Libraries were prepared by the University of Colorado Genomics Shared Resource using low-input ribosomal-depleted total RNA library construction service (SMARTer Stranded Total RNA-Seq Kit v2 – Pico Input Mammalian, Takara Bio) and sequenced paired-end 150 bp on an Illumina NovaSeq X, targeting ≥50 million reads per sample. Samples were sequenced across two batches. Samples with genome coverage below 15% were excluded from downstream analysis, resulting in a total of 12 IP samples (N=3 per sex and treatment group) and 11 Input samples for subsequent data analysis. One Input sample (male saline group) was excluded due to QC result.

#### Immunohistochemistry

Mice were deeply anesthetized with isoflurane and underwent trans-cardiac perfusion of cold 1x Phosphate-Buffered Saline (PBS) followed by cold 4% paraformaldehyde (PFA) in 1x PBS. LS-DRG (L1, L2, L6, S1, and S2) were bilaterally collected. Isolated DRG were post-fixed in 4% PFA overnight, cryoprotected with 30% sucrose for 7 days at 4°C, and frozen in Optimal Cutting Temperature (OCT) compound using a dry ice/ethanol bath. Samples were stored at −80°C until sectioning. DRG samples were sectioned using a Leica cryostat (CM1520) at a thickness of 20 µm and stored at −20°C prior to immunostaining.

DRG tissue sections were permeabilized with 0.2% Triton x-100, blocked for 1hr in 10% donkey serum (containing 0.2% Triton), and incubated overnight with primary antibody diluted in 10% donkey serum containing 0.2% Triton x-100. Rabbit anti-brain lipid-binding protein (BLBP) (Millipore ABN14, 1:500) or rabbit anti-myelin protein zero (MPZ) (Cell Signaling Technology 57518, 1:250) was used to label SGCs or Schwann cells, respectively. After removing excess primary antibodies, tissue sections were then incubated with donkey anti-rabbit secondary antibody (Invitrogen, A-31573,1:500) for 3 hours. DAPI (Invitrogen, D1306, 1 µM) was used to label cell nuclei. Slides were then washed with 1x PBS and allowed to air-dry before coverslipping.

#### Confocal imaging

Images were taken using a Zeiss LSM 780 Confocal Microscope. GFP signal was imaged using a 488nm laser at 5% power; OFP was imaged with a 561 nm laser with 3% laser power, and BLBP or MPZ signals were imaged with a 633nm laser at 3% laser power. DAPI was imaged using a 405 nm laser at 1% laser power. DRG images were taken using 20x and 40x objectives and randomized.

#### Cytokine study

A separate cohort of C57BL/6J mice of both sexes was randomly assigned to no instillation (naïve), saline instillation or VEGF instillation (N = 12 per group; 6 males and 6 females). LS-DRG were harvested from instilled animals on day 16 after the first instillation, and from naïve animals and tissue was homogenized in PBS containing protease inhibitors using a Precellys homogenizer in a Precellys CK Mix tubes (Cat. No.: P000918-LYSK0-A, Bertin Instruments) in 10-second intervals at medium speed, repeated 8 times, in the cold room. The lysate was centrifuged for 5 minutes at 12,000×g at 4°C. Total protein concentration in DRG tissue lysates was determined by bicinchoninic acid (BCA) assay. The cytokine expression was then assayed using a Bio-Plex Pro Mouse Cytokine Group I 23-Plex Assay (Catalog #: M60009RDPD; Bio-Rad Laboratories, Hercules, CA) following the manufacturer’s protocol for tissue lysates. In brief, 25 µg of protein was loaded per well. Standards were reconstituted in the kit-provided standard diluent per manufacturer’s instructions. Magnetic beads were incubated with samples and standards, washed, and detected per kit protocol. Bead fluorescence was acquired using a Bio-Plex® 200 Multiplex Suspension Array System (Reader Serial Number: LX10018157421). Mean fluorescence intensity (MFI) was converted to concentration (pg/mL) using a 5-parameter logistic standard curve fit per analyte. Samples were analyzed in technical duplicates. Concentrations were calculated as the means of both replicates when both were above the lower limit of quantification (LLOQ), the single detectable value when only one replicate exceeded the LLOQ and recorded as not detected when both replicates were below the LLOQ. Analytes undetectable in all samples (Supplemental Table S4) were excluded from further analysis.

### QUANTIFICATION AND STATISTICAL ANALYSIS

#### Single-cell and single-nucleus atlas reanalysis

Raw RNA sequencing data from publicly available mouse DRG snRNA-seq (ref. 30; GSE201653) and scRNA-seq (ref. 14; GSE174430) datasets were downloaded and processed independently in Partek Flow (version 12.10.0, Partek Inc., an Illumina company, St. Louis, MO, USA). Reads were aligned to GRCm39 (Ensembl release 105) and quantified with Cell Ranger (v9.0.0), with intronic reads included. Nuclei passing barcode QC were normalized (log2[CPM + 1]), integrated across samples with Harmony, and clustered with the Leiden algorithm on the top 2,000 variable genes. The SGC cluster was identified by enrichment of canonical SGC markers (Apoe, Kcnj10, Fabp7, Sox10 and Megf10). Raw counts from all SGC nuclei or cells in each sample were summed into pseudobulk profiles and normalized to counts per million. For the scRNA-seq reference, only naïve control cells from the Cell_SNI dataset (GEO: GSE174430; Jager et al., 2022) were included; these samples are male only, so no averaging across sexes was performed for this dataset. For the snRNA-seq dataset (GSE201653), which comprises one male and one female sample, pseudobulk profiles were averaged across the two sexes.

#### SGC and non-SGC marker panels

For each atlas, SGC-enriched genes were defined as FDR<0.05 and ≥2-fold higher expression in the SGC cluster than in other clusters. For display in Figure 1E, a stricter specificity filter (least-squares mean >20 in the SGC cluster, <5 in all other clusters, FDR<0.01) was applied, and the top 25 genes per atlas were selected. Non-SGC marker panels (five genes each for fibroblasts, neurons, Schwann cells, endothelial cells, pericytes and immune cells) were taken from the atlas. Per-gene enrichment was tested with two-sided paired Wilcoxon signed-rank tests of log₂(IP/Input) against zero with Benjamini–Hochberg correction within each panel, and SGC versus non-SGC panel scores were compared using a two-sided paired Wilcoxon signed-rank test across the 11 matched pairs.

#### Translatome CPM quantification

Raw count matrices from 11 matched TRAP-IP and bulk DRG Input samples were used for gene-level quantification. Library-size normalization was performed by scaling raw counts to counts per million (CPM) using sample-specific library sizes. The mean IP CPM and mean Input CPM were calculated per gene across all 11 pairs. A curated panel of gap junction and pannexin transcripts, GPCR signaling genes, and housekeeping reference genes was selected for comparative visualization across the three data sources. Reads were aligned to GRCm39 with STAR v2.7.8a (default parameters) and quantified with HTSeq (v0.11.0, union mode, unstranded) against Ensembl release 105 in Partek Flow (v12.10.0), retaining uniquely mapped reads only (MAPQ = 255). Multimapping reads were not counted. Genes were ordered by descending mean IP CPM within each category. All plots were generated in R using ggplot2 and patchwork.

#### GPCR repertoire analysis

GPCR-type entries (413) were taken from the IUPHAR/BPS Guide to Pharmacology target list (v2026.2)^46^ and mapped to 355 distinct mouse genes. After excluding IUPHAR-designated non-GPCR 7TM proteins, Adgrd2-ps, Gpr31c, chemosensory receptors and genes absent from the translatome count matrix, 343 receptors were evaluated using all 11 TRAP-IP/Input pairs. Presumed chemosensory receptors that did not meet the sorted-SGC expression criterion (2,039 genes) served as negative controls, and the detection floor was set at the 95th percentile of their mean TRAP-IP CPM. Receptors above the floor were retained if they had FPKM >1 in at least three of four replicates of published bulk RNA-seq from FACS-sorted naïve SGCs and were excluded if cell-weighted non-SGC expression was ≥2.5-fold higher than SGC expression in each of the two DRG atlases. TRAP-IP-to-Input ratios were computed within each animal as log₂[(IP CPM + 1)/(Input CPM + 1)] and averaged across the 11 pairs. As a sensitivity analysis, the full selection pipeline was repeated using only the five saline pairs.

CPM is not length-normalized, so between-gene comparisons favor longer transcripts^49^; the within-animal IP/Input ratio is unaffected by transcript length.

#### Differential expression analysis between sexes

Differential expression analysis was performed using DESeq2 (v1.50.2; Love, Huber & Anders, 2014) in R (v4.5.3). A paired DESeq2 model (∼ pair + fraction) was fit to matched TRAP-IP –Input and genes were called TRAP-enriched at BH-adjusted p<0.05 and log₂FC(IP/Input)>0. Sex differences were modelled using DESeq2 with sequencing batch as a covariate (∼ batch + sex), and significance was assessed using Wald tests. DEGs were defined as nominal p<0.05 and fold change >1.5. Genes were classified as autosomal protein-coding, sex-linked (X/Y), or non-coding based on Ensembl annotations. DEGs detected in the TRAP-IP but not in matched Input fractions were designated translatome-specific. Figures were generated in R (v4.5.3) using ggplot2 (v4.0.3), ggrepel (v0.9.8) and patchwork (v1.3.2).

#### Differential expression analysis between VEGF and saline instillations

Translational responses to VEGF were assessed by DESeq2 (design: ∼batch + sex + treatment; contrast VEGF vs. Saline), restricted to autosomal protein-coding. Genes were classified as VEGF-up- or down-regulated using a nominal p<0.05 and >1.5-fold change. Ingenuity Pathway Analysis (IPA) was performed on the treatment-effect gene list (p<0.05) to identify candidate upstream regulators and enriched canonical pathways. Eif4e was identified as the only predicted activated translational regulator among the differentially expressed genes. Representative functional gene sets were selected from IPA-enriched pathways (cytokine/immune signaling) or curated from differentially expressed genes representing major biological processes identified in the EIF4E regulatory network, including vesicle/adaptor trafficking and mitochondrial function. Variance-stabilized (VST) expression values were z-scored for each gene across all 12 individual TRAP-IP samples. The resulting z-scores were then averaged within the Saline (N=6) and VEGF (N=6) groups for heatmap visualization.

#### Candidate SGC functional gene panel

To examine known SGC biosignaling alteration in response to VEGF instillations, 15 functional categories and their constituent genes were selected using independent DRG scRNA-seq and snRNA-seq datasets processed in Partek Flow. Genes were retained if their mean SGC-cluster expression was at or above the 33rd percentile of all genes’ SGC-cluster means in either dataset. Mean expression across all non-SGC clusters also had to be less than 2.5-fold than the SGC mean in either dataset. Both atlases are independent of the TRAP-IP counts and of treatment, so the restriction is not circular. The study-wide threshold was nominal two-sided p<0.05 together with | log₂ fold-change| >log₂(1.5). Benjamini–Hochberg correction was applied across the 133 unique genes of the prespecified panel. Counts were modeled using DESeq2. Sex-stratified treatment contrasts used the design ∼ batch + treatment; the combined treatment contrast used ∼ batch + sex + treatment; and sex dependence of the treatment effect was tested using ∼ batch + sex + treatment + sex:treatment. For each category in the male, female, and combined treatment models, the change detectable with approximately 80% power at nominal two-sided α=0.05 was estimated as 2.8 times the median standard error. Across categories, detectable changes were 1.54-to 2.51-fold in the combined model and 2.01-to 2.85-fold in the sex-specific models. No pathway-level test was applied.

#### Analysis of hormonal influence on the SGC translatome

Detection of Ar, Esr1 and Esr2 was evaluated in independent DRG scRNA-seq and snRNA-seq datasets using Partek Flow feature-statistics exports. SGC clusters were pooled within each platform, and neurons were used as a reference population. Detection was calculated as the percentage of cells or nuclei with nonzero counts. Because biological replicate-level detection data were unavailable, the analysis was descriptive, and no inferential statistical tests were performed.

Female and male VEGF vs saline TRAP gene lists were analyzed separately using IPA. Genes with p<0.05 together with |log₂ fold-change| >log₂(1.5) were included. IPA used known biological relationships to show how these genes may connect with the androgen receptor (AR) and estrogen receptor-α (ESR1) signaling. AR and ESR1 were included as reference nodes and were not differentially translated genes; therefore, their presence does not demonstrate receptor expression or activation in SGCs.

#### Bulk RNA-seq of DRG cytokine transcripts

Bulk LS-DRG RNA-seq from a previously published cohort (Yesupatham et al., 2026; GEO: GSE294967) was reanalyzed. The dataset contains 16 animals, with four animals per sex × treatment group. Reads were not realigned for the present study; gene-level counts generated in the original publication were used directly. As reported there, reads were aligned with STAR v2.7.8a and quantified with HTSeq against the GRCm39 (mm39) genome with Ensembl annotation in Partek Flow. Each Bio-Plex analyte was mapped to its encoding mouse gene. A transcript was considered detected when its mean normalized count was ≥10 with at least 3 samples at raw count ≥5. VEGF versus saline was tested by DESeq2 (∼sex + treatment) at nominal p < 0.05; genes with zero counts in all samples are dropped by DESeq2 and could not be tested. Aligned reads were inspected in a genome browser for every analyte meeting the count-based detection criterion; analytes whose counts arose from non-exonic read stacks or from internal priming artifacts were not considered reliably detected, regardless of count magnitude. Values are plotted per animal as log10(normalized count + 1).

Group values are reported as medians. Significance was assessed at a nominal threshold of p<0.05 and by Benjamini–Hochberg correction across the tested analytes.

#### Behavior, imaging and cytokine statistics

GraphPad Prism (version 10) was used to analyze the data and generate figures. The withdrawal frequencies were analyzed using two-way ANOVA with sex/genotype/treatment and filament forces as between subjects’ factors. All data are expressed as the mean ± SEM.

GFP+, OFP+, BLBP+, and MPZ+ cells were counted in Fiji (ImageJ, v2.14.0) from each image (3∼5 DRG image per animal; Cre+ with Tamoxifen injections: n=11 images/3 animals; Cre+ with corn oil: n=3 images/1 animal). Figures were analyzed and generated using GraphPad Prism (version 10). All data are expressed as the mean ± SEM.

Per-sample concentrations (pg/mL) were grouped by cytokine, sex, and treatment (Naïve, Saline, VEGF). Figures were analyzed and generated using GraphPad Prism (version 10). For each analyte–sex–treatment combination with ≥2 animals, a boxplot (median, interquartile range) was overlaid with individual data points; groups with fewer than 2 animals are shown as individual points. Concentrations were log10-transformed and analyzed by ordinary two-way ANOVA with treatment, sex and their interaction, followed by Tukey’s multiple comparisons among treatments within each sex. Values below the LLOQ were excluded from statistical analysis and are displayed at LLOQ/2 for visualization only.

Statistical details for each experiment (test, n and what n represents, and definition of center and dispersion) are provided in the figure legends.

## Acknowledgements

This research project is supported by NIDDK R01DK129260 (Xie). AG’s internship was supported by Colorado Research Experiences (NINDS 1R25NS130620). The *L10a-GFP* mice was a generous gift from the late Dr. Marc G. Caron (Duke University). The *Sox10-CreER^T2^* mice was a generous gift from Dr. Brian D. Gulbransen (University of Nevada, Reno) with the permission from Dr. Vassilis Pachnis (The Francis Crick Institute, UK) who developed the mouse line. The authors would like to thank the University of Colorado Genomics Shared Resources (supported by the Cancer Center Support Grant P30CA046934) for assisting with cDNA library preparation and RNA sequencing, Dr. Jeremy Rahkola at the Veterans Affairs Research Core Facility (VA Eastern Colorado Health Care System, Aurora, CO) for assisting with cytokine profiling, and the Advanced Light Microscopy Core (partially supported by the Diabetes Research Center Grant P30 DK116073). Special thanks to Dr. Sushil Kumar for his helpful suggestions on data analysis and statistics, and Ms. Angelin Tisha for helping with the R coding and data visualization. Lastly, we would like to thank the authors of the papers from which we used sequencing data for their exemplary transparency in sharing the details of their work, including a single-nucleus RNA-seq atlas GSE201653, a single-cell RNA-seq atlas GSE174430, and bulk RNA-seq of FACS-sorted satellite glial cells GSE120284.

## Author contributions

AXX obtained the funding for the study. SKY and AXX designed the study. SKY maintained the mouse colony and conducted most of the experiments. AXX and AG performed Von Frey testing and immunohistology, respectively. All authors analyzed data and wrote the paper.

## Declaration of generative AI and AI-assisted technologies in the manuscript preparation process

During the preparation of this work, the authors used Claude (Anthropic) to write R code for data visualization and to improve the language and readability of the manuscript. After using this tool, the authors reviewed and edited the content as needed and take full responsibility for the content of the published article.

---

**Supplemental Fig. S1.**
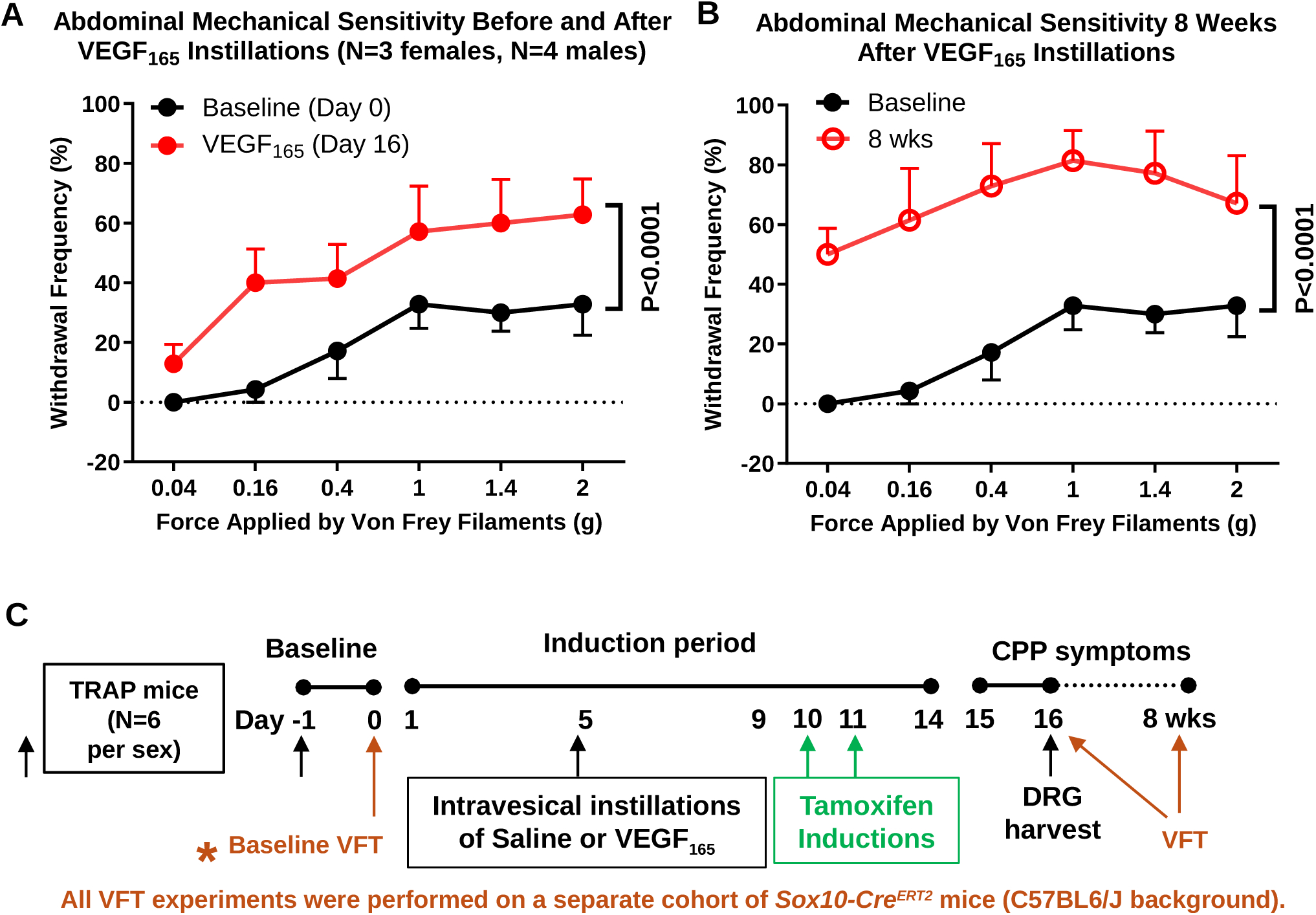
Intravesical instillations of VEGF_165_ and VEGFR1 agonist induces pelvic mechanical hypersensitivity. (A) Experimental paradigm and timeline. (B) VEGF_165_ instillation over 9 days increase lower abdomen withdrawal frequencies to Von Frey filaments on day 16. (C) VEGF165-induced pelvic mechanical hypersensitivity lasts at least 8 weeks. (D-E) VEGF_B_, the VEGFR1 agonist, but not VEGF_E_, the VEGFR2 agonist, resulted in increases in pelvic mechanical sensitivity on day 16 comparable to that observed following VEGF_165_. (F) VEGF_E_ induces visceral hypersensitivity 5-6 weeks after intravesical instillations.

**Supplemental Fig. S2.**
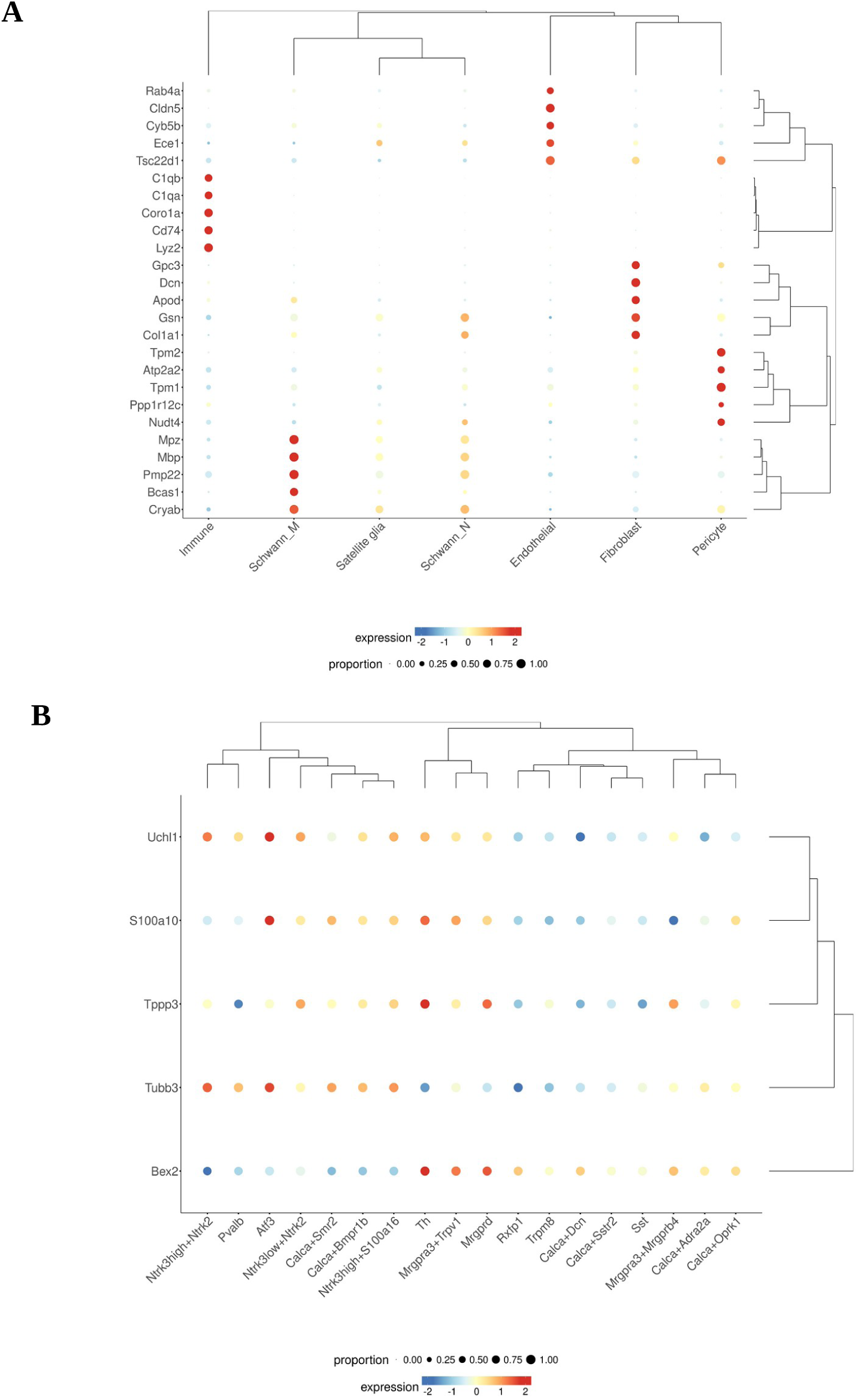
Cell-type specificity of non-SGC and neuronal marker genes in the mouse naïve DRG single-cell atlas. Bubble plots showing expression of selected marker genes across DRG cell types from the harmonized mouse naïve DRG sc/snRNA-seq atlas (Bhuiyan et al., 2024, *Science Advances* 10, eadj9173), visualized using the PainSeq portal. (A) Non-SGC marker genes across non-neuronal DRG cell clusters including immune, Schwann, endothelial, fibroblast, pericyte, and satellite glial cells. Marker panels showed strong enrichment in their respective target cell types with low expression in satellite glial cells, supporting their use as non-SGC depletion markers. (B) Neuronal marker genes (*Uchl1, S100a10, Tppp3, Tubb3,* and *Bex2*) across mouse DRG neuronal subtypes. All markers showed broad neuronal expression across multiple neuronal populations, confirming their suitability as pan-neuronal markers. Dot color represents scaled normalized expression level, and dot size indicates the proportion of cells expressing each gene.

**Supplemental Fig. S3.**
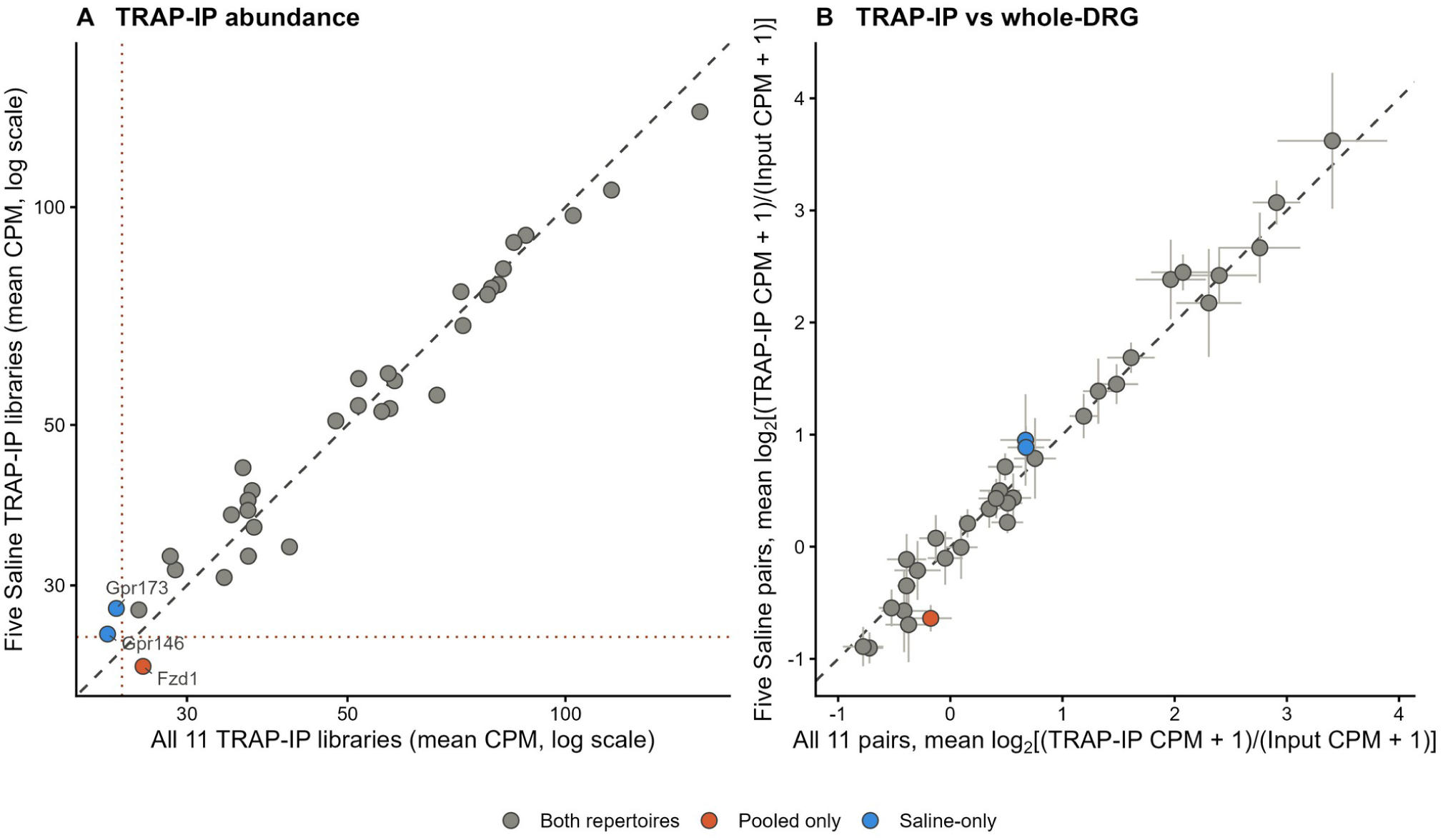
Saline-only sensitivity analysis of the GPCR repertoire. The complete selection pipeline was repeated using only the five saline-treated animals. This identified 33 receptors: 31 were shared with the 32-receptor repertoire from the full analysis, *Fzd1* was retained only in the full analysis, and *Gpr146* and *Gpr173* only in the saline analysis. All three were close to the detection floor. Grey points are receptors shared by both analyses, orange the receptor retained only in the full analysis (*Fzd1*), and blue those retained only in the saline analysis (*Gpr146*, *Gpr173*).. (A) Mean TRAP-IP abundance across all eleven animals versus the five saline-treated animals. The dashed line shows equality, and the dotted lines mark the corresponding detection floors of 24.4 and 25.5 CPM. Labelled receptors changed repertoire membership. Across the 31 shared receptors, Pearson r=0.980 on log₁₀ CPM. (B) Mean TRAP-IP-to-input ratios from all eleven matched pairs versus the five saline pairs. The dashed line shows equality. Error bars show the standard error across pairs on each axis and were wider in the saline-only analysis because fewer animals were included (median inflation-fold).

**Supplemental Fig. S4.**
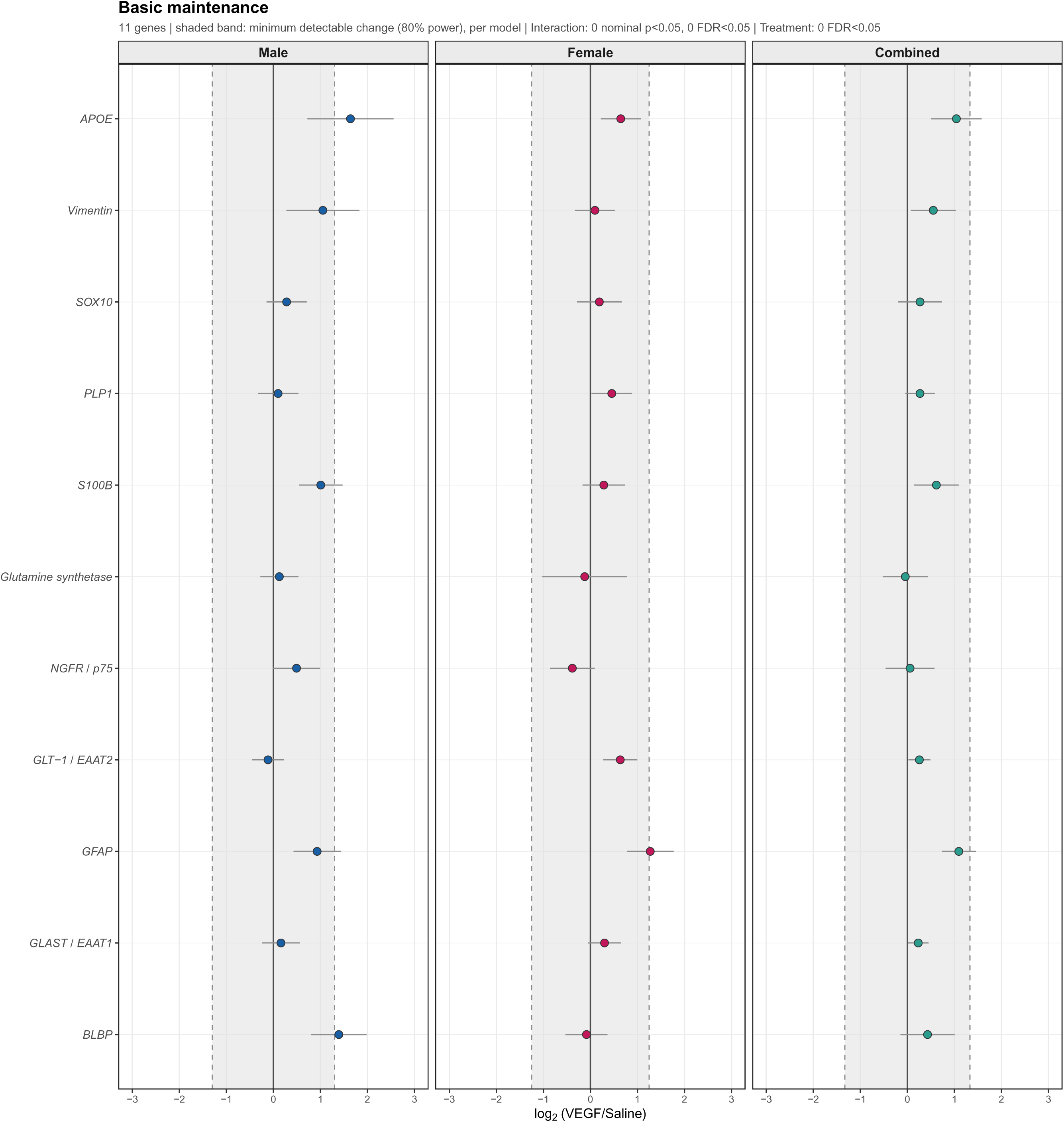
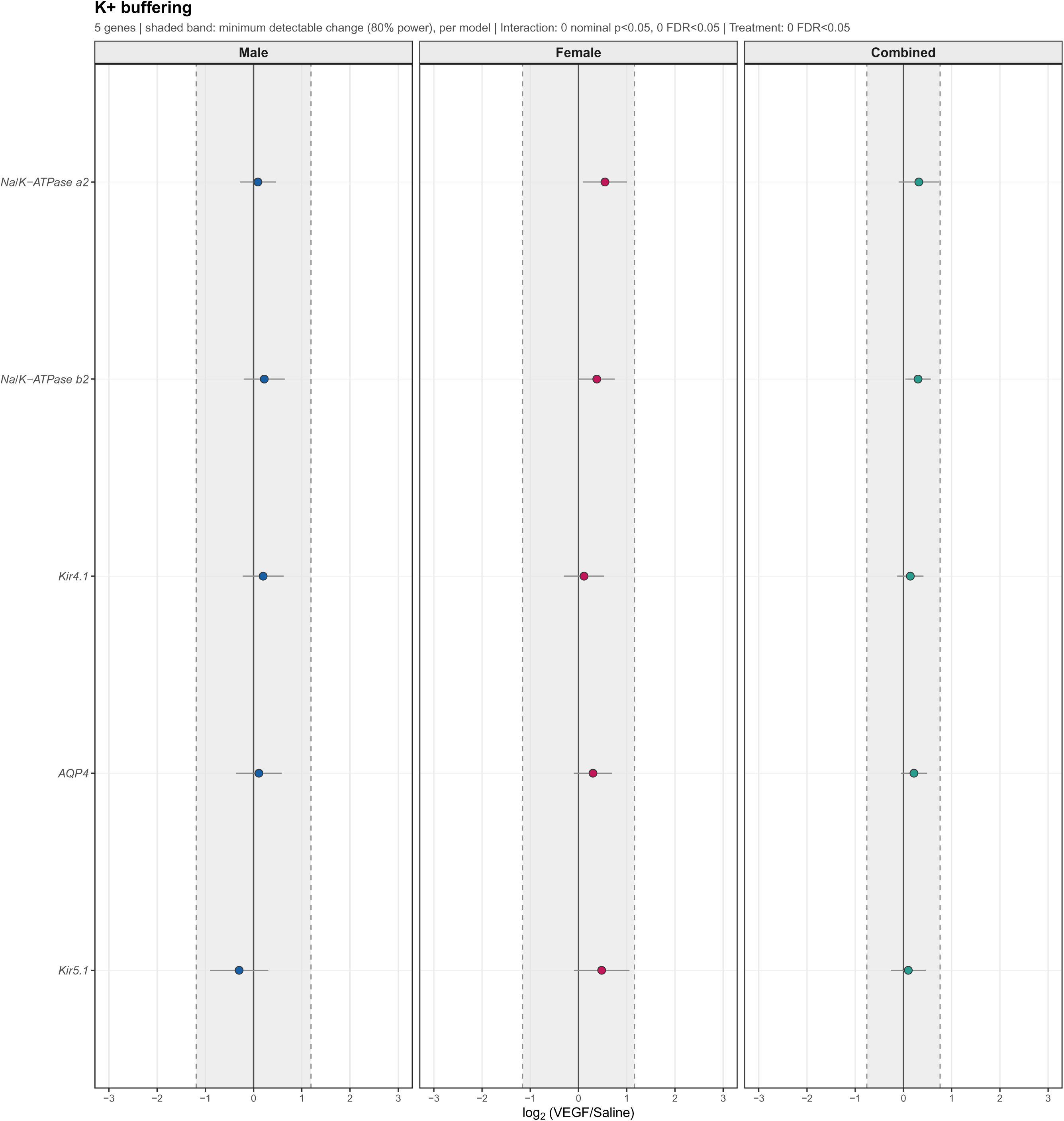
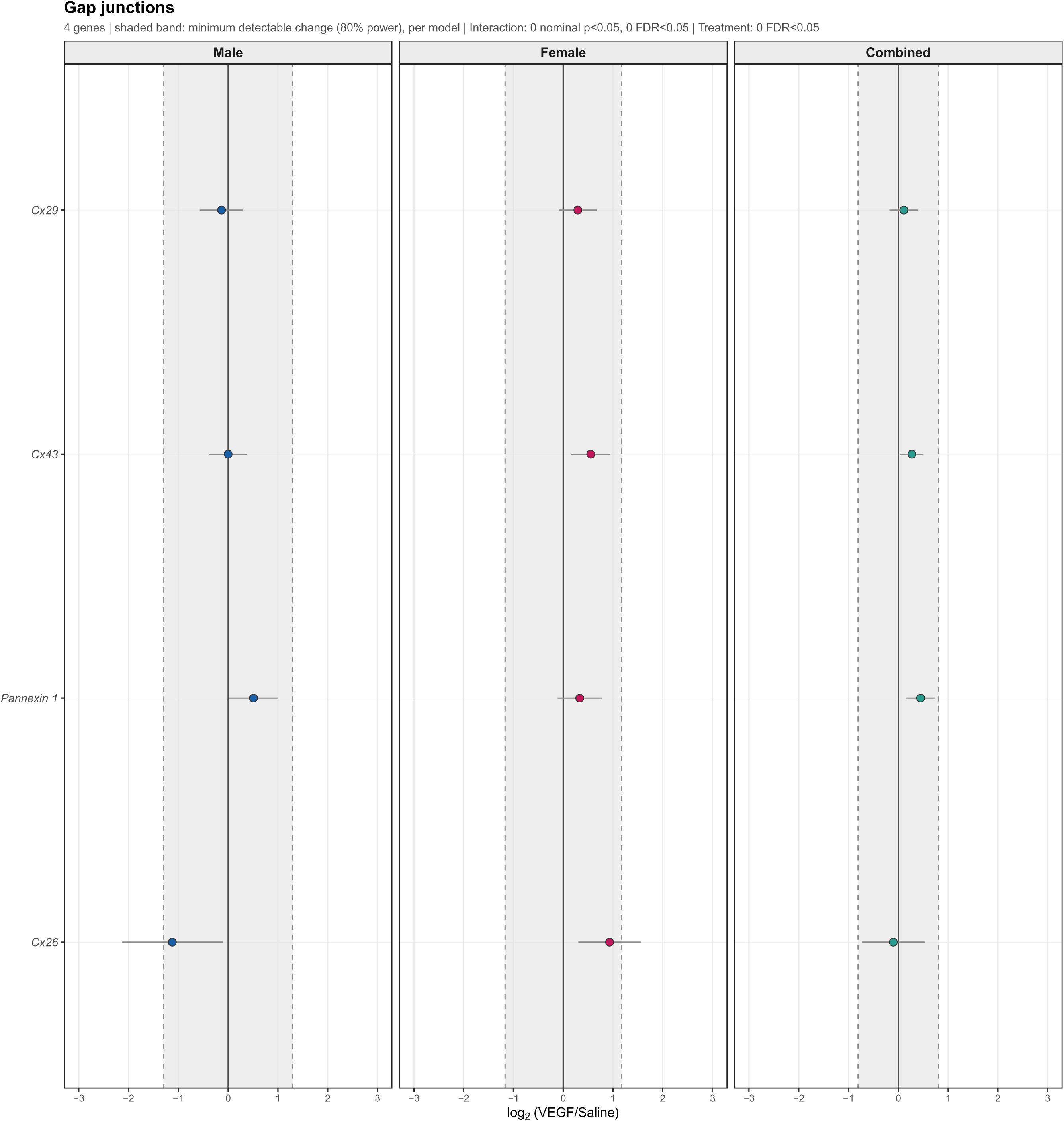
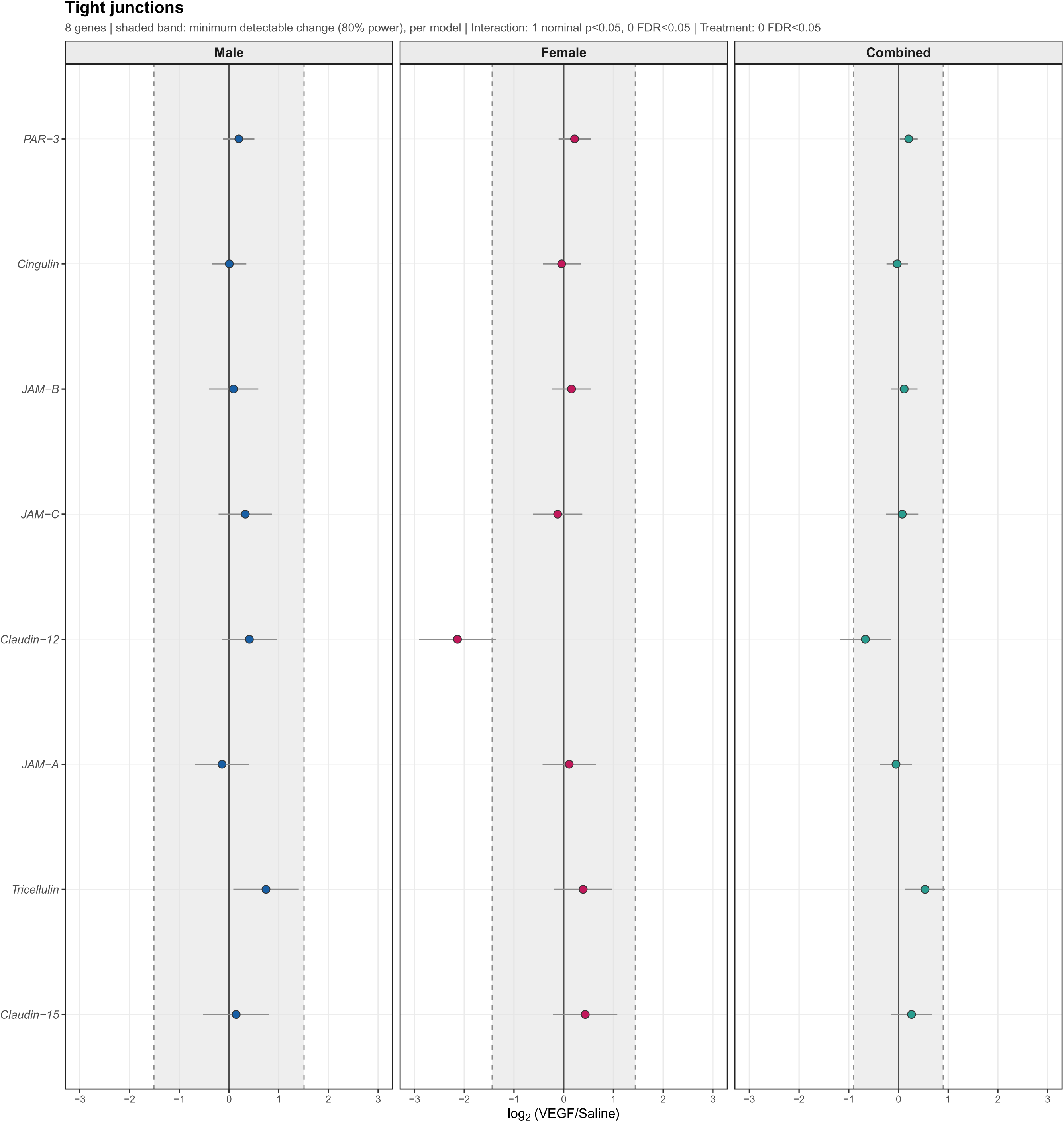
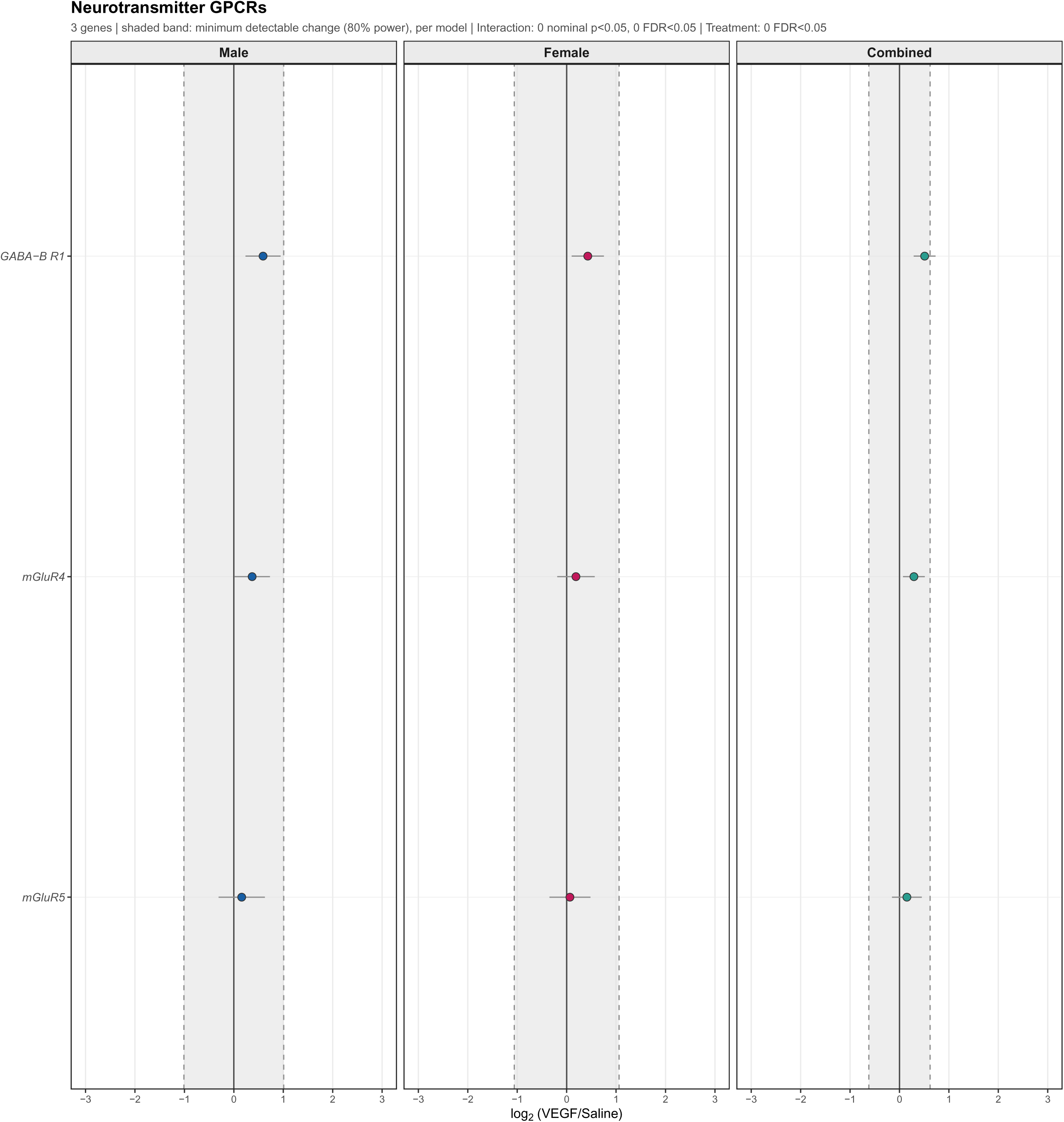
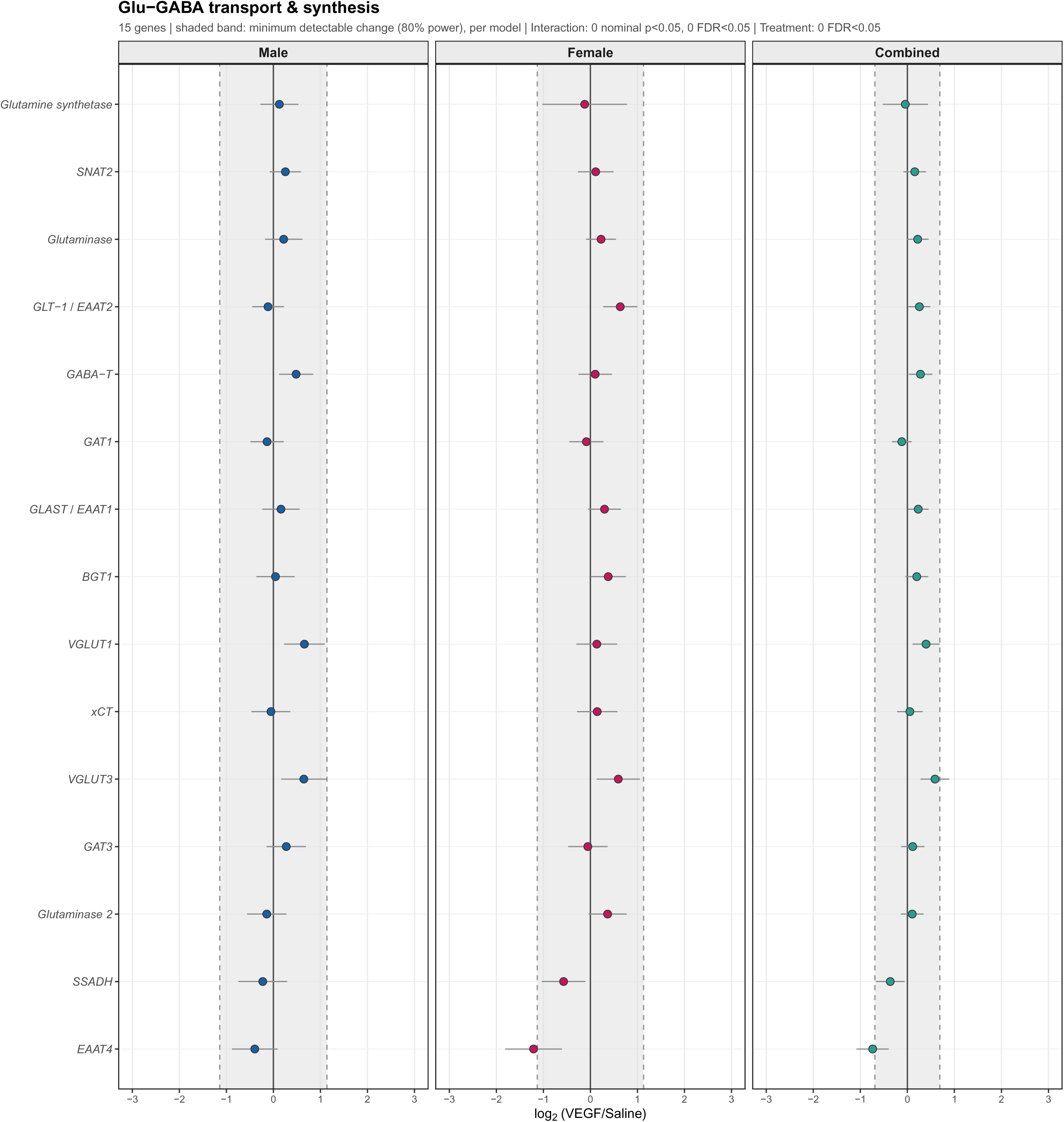
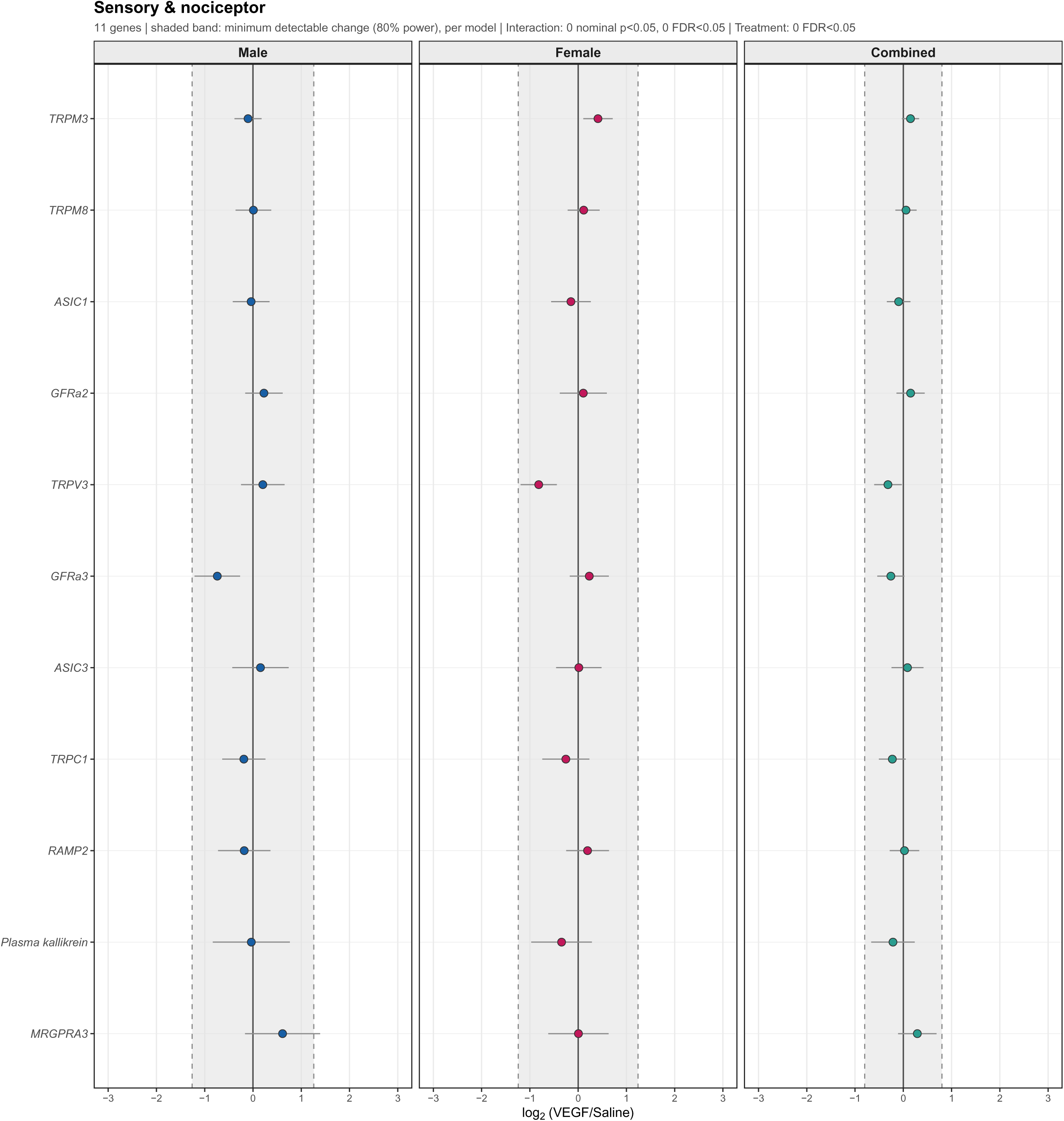
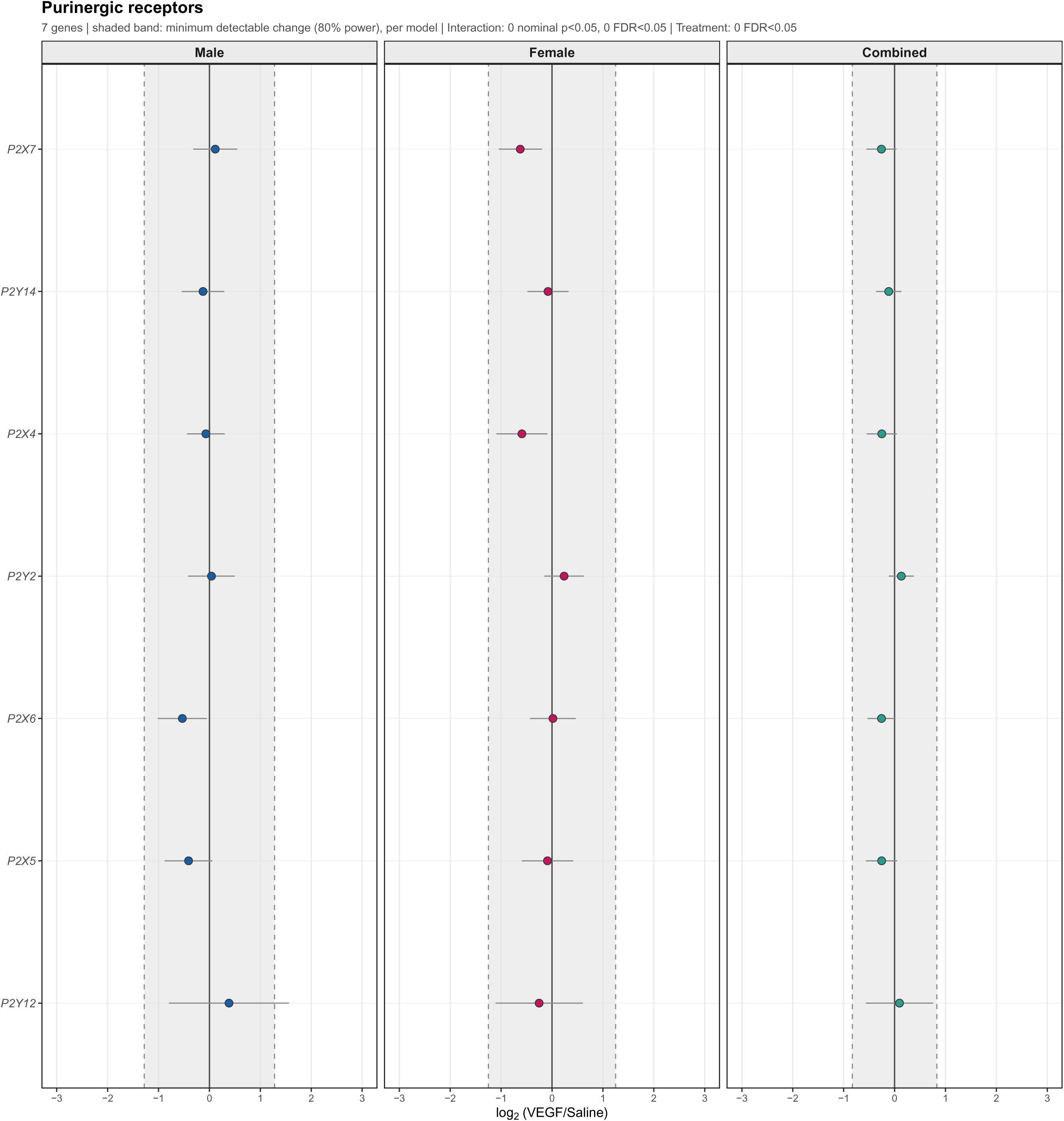
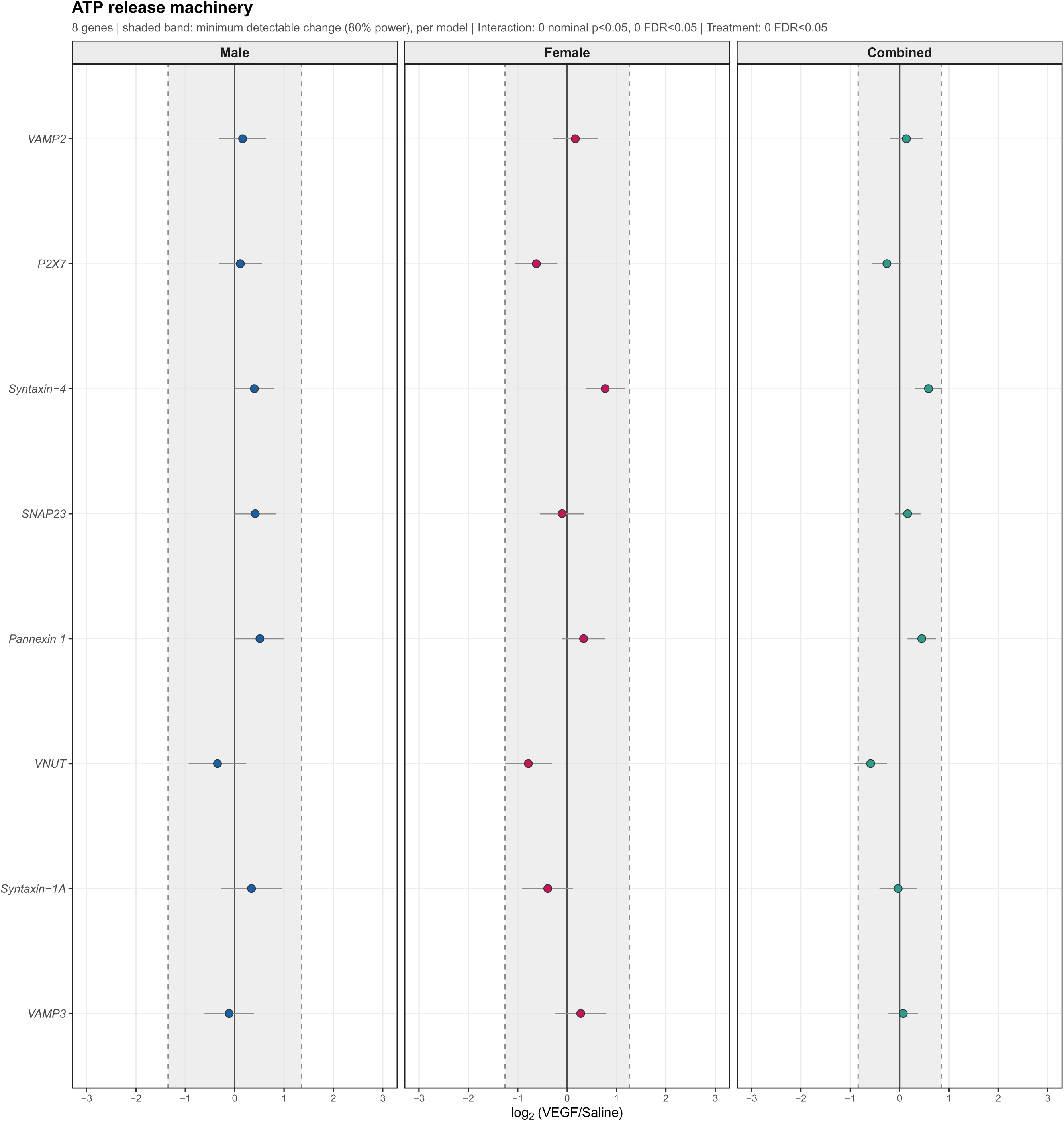
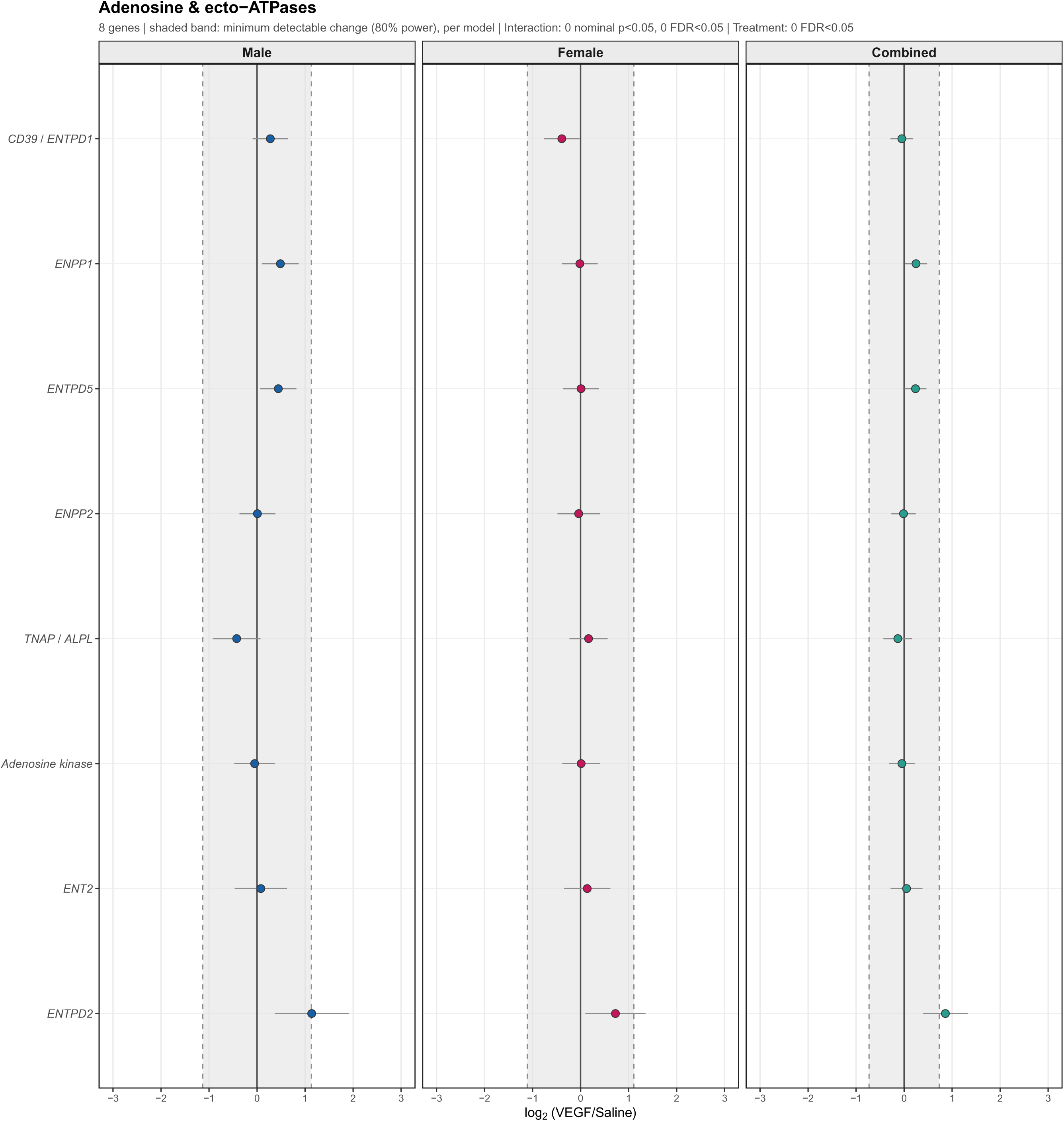
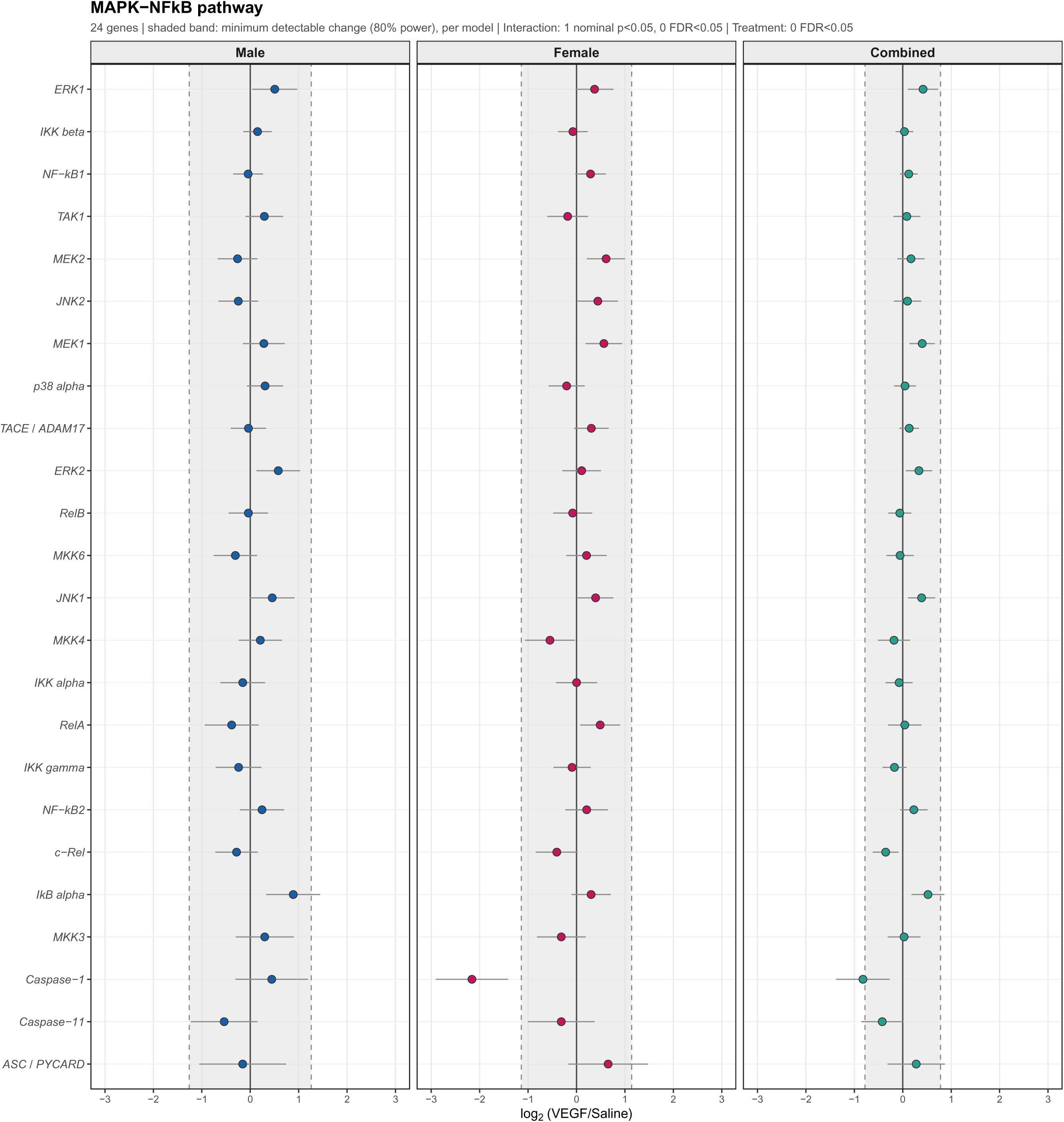
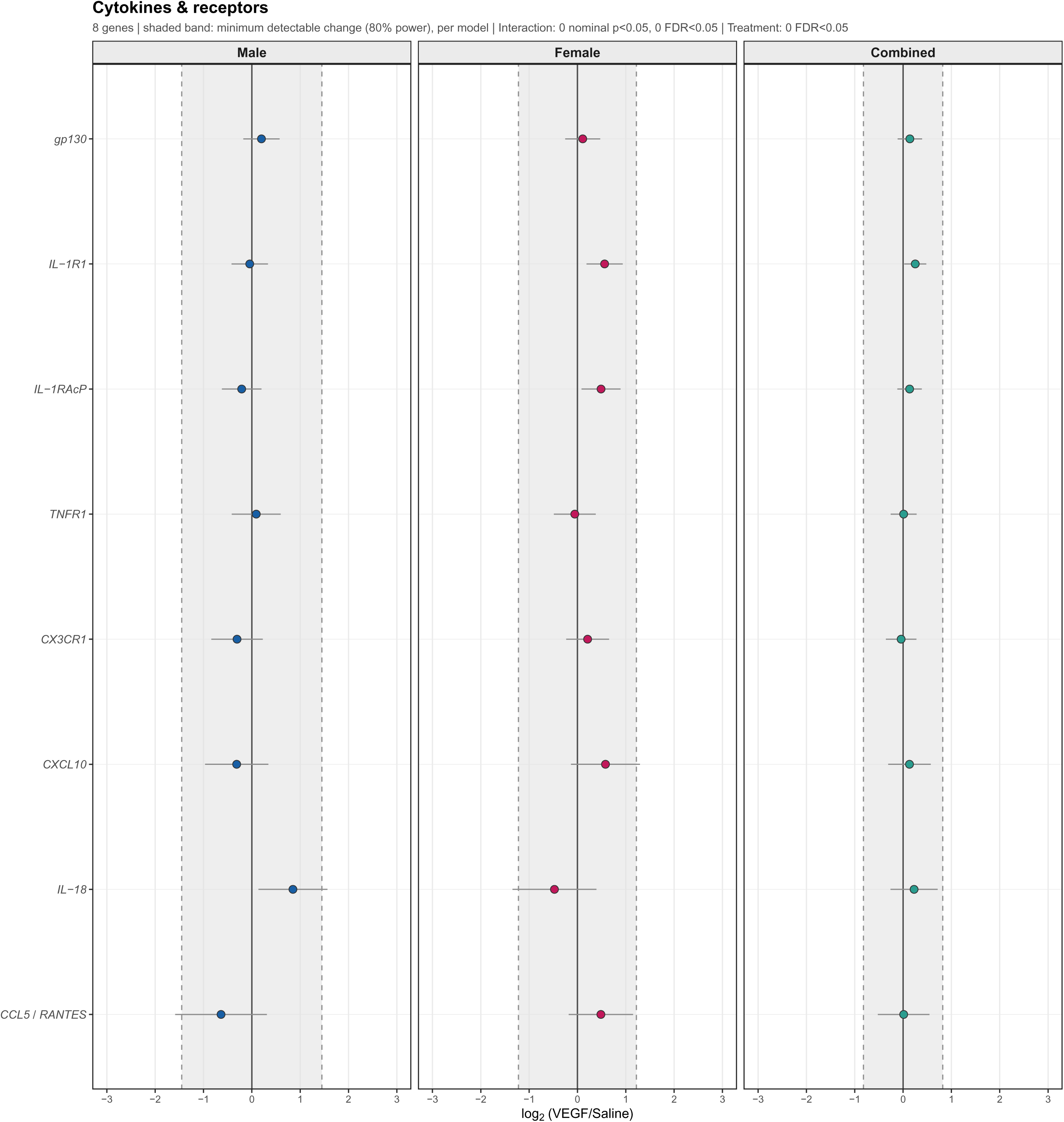
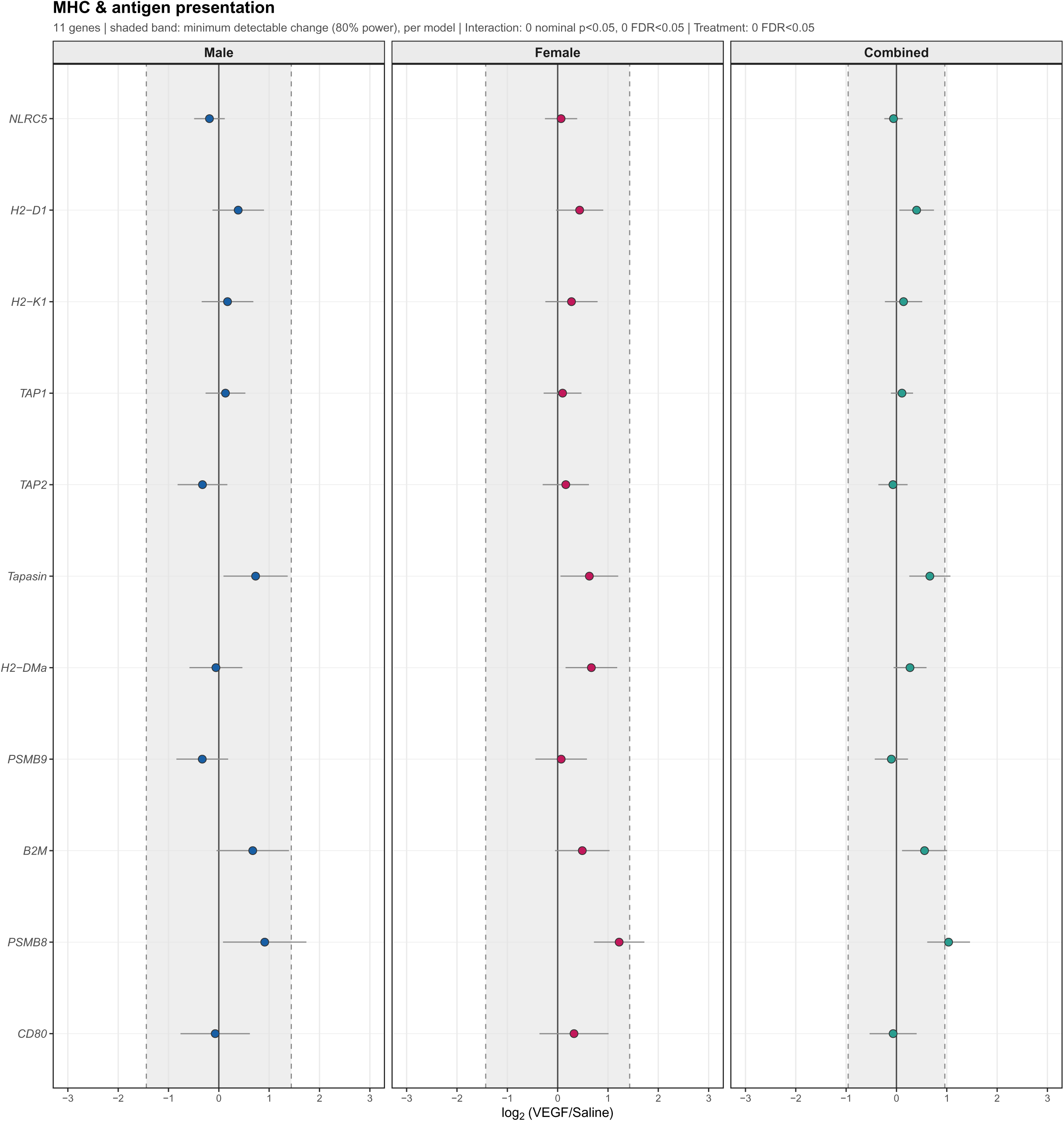
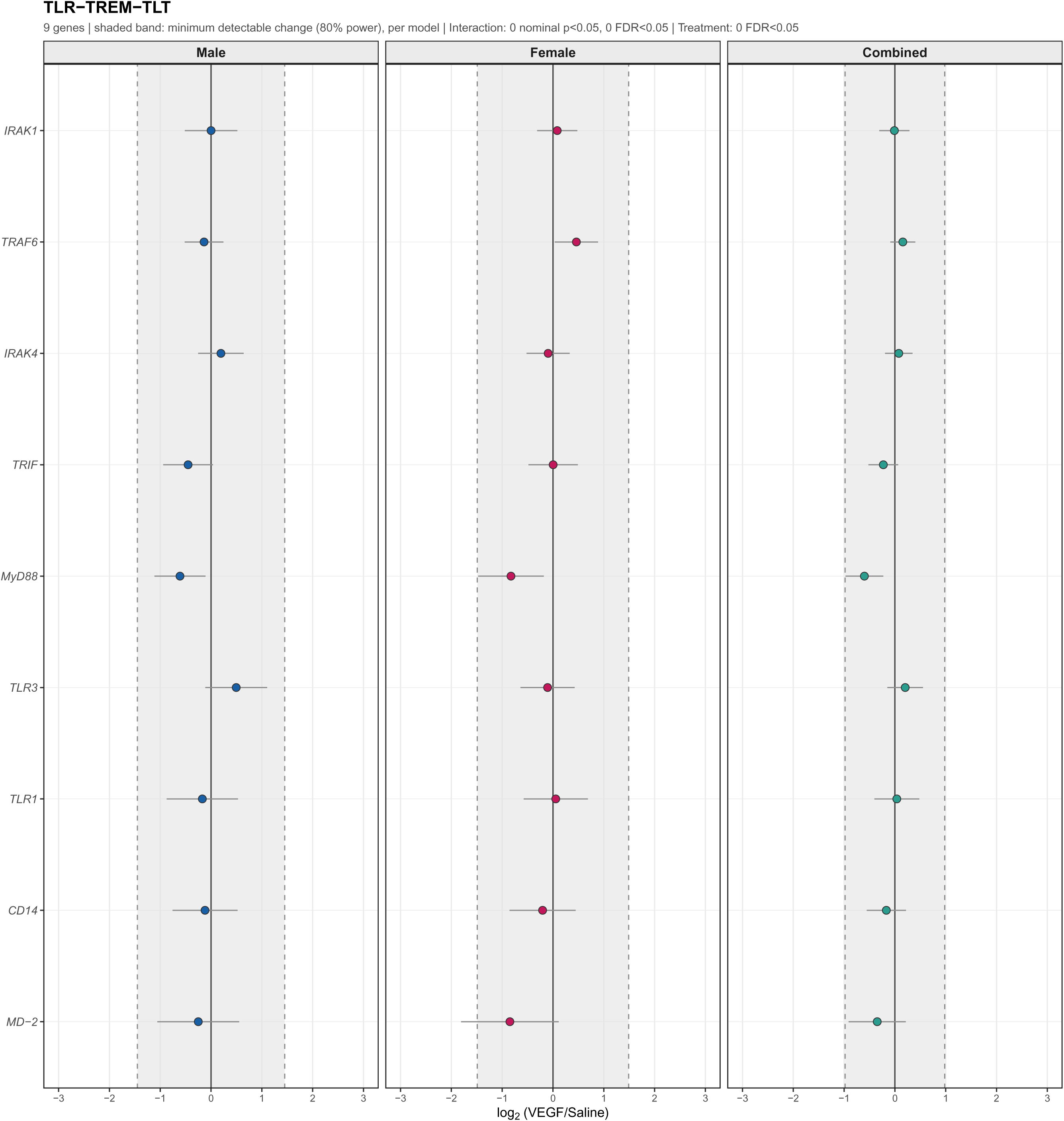
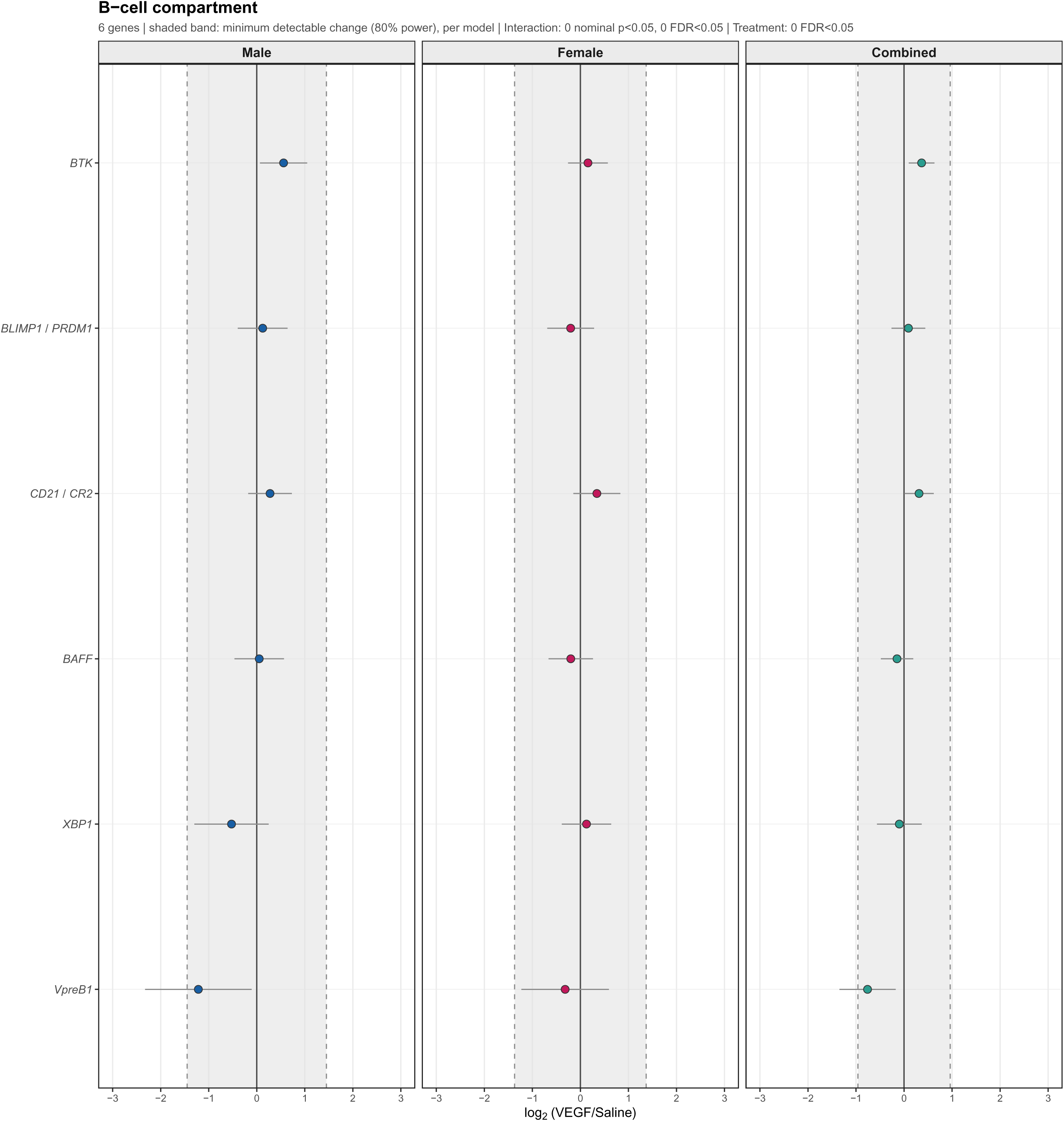
VEGF-associated changes in an a priori SGC functional gene panel. Plots show VEGF-associated log₂ fold changes for 133 genes across 15 functional categories. Genes were retained using independent DRG atlases (SGC expression ≥33rd percentile and non-SGC/SGC ratio <2.5 in either atlas). Points show estimated log₂ fold changes (VEGF/saline), with error bars indicating ±SE, for males, females and both sexes combined. Grey bands mark the approximate minimum changes detectable with 80% power. Genes are ordered by mean IP CPM.

